# The ciliary MBO2 complex targets assembly of inner arm dynein *b* and reveals additional doublet microtubule asymmetries

**DOI:** 10.1101/2023.07.31.551375

**Authors:** Gang Fu, Katherine Augspurger, Jason Sakizadeh, Jaimee Reck, Raqual Bower, Douglas Tritschler, Long Gui, Daniela Nicastro, Mary E. Porter

## Abstract

Ciliary motility requires the spatiotemporal coordination of multiple dynein motors by regulatory complexes located within the 96 nm axoneme repeat. Many organisms can alter ciliary waveforms in response to internal or external stimuli, but little is known about the specific polypeptides and structural organization of complexes that regulate waveforms. In *Chlamydomonas*, several mutations convert the ciliary waveform from an asymmetric, ciliary-type stroke to a symmetric, flagellar-type stroke. Some of these mutations alter subunits located at the inner junction of the doublet microtubule and others alter interactions between the dynein arms and the radial spokes. These and other axonemal substructures are interconnected by a network of poorly characterized proteins. Here we re-analyze several motility mutants (*mbo*, *fap57*, *pf12/pacrg*) to identify new components in this network. The *mbo* (*m*ove *b*ackwards *o*nly) mutants are unable to swim forwards with an asymmetric waveform. Proteomics identified more than 19 polypeptides that are missing or reduced in *mbo* mutants, including one inner dynein arm, IDA *b*. Several MBO2-associated proteins are also altered in *fap57* and *pf12/parcg* mutants, suggesting overlapping networks. Two subunits are highly conserved, coiled coil proteins found in other species with motile cilia and others contain potential signaling domains. Cryo-electron tomography and epitope tagging revealed that the MBO2 complex is found on specific doublet microtubules and forms a large, L-shaped structure that contacts the base of IDA *b* that interconnects multiple dynein regulatory complexes and varies in a doublet microtubule specific fashion.

## Introduction

Eukaryotic cilia and flagella are microtubule-based extensions of the cell surface important for cell motility and cell signaling. Defects in cilia or flagella have been linked to a complex group of diseases collectively termed ciliopathies (reviewed in Reiter and Leroux, 2017). In vertebrate embryos, motility is required for the rotary movement of nodal cilia that determines the left-right body axis (reviewed in Babu and Roy, 2023). In adults, motility is essential for the circulation of cerebrospinal fluid, the clearance of particles and mucus in the respiratory tract, and sperm motility. Defects in the assembly, beating, or signaling of motile cilia have been linked to situs inversus, hydrocephalus, curvature of the spine, chronic respiratory disease, and infertility (Mitchison and Valente, 2017). Given the complex structure of cilia and the large number of genes involved in their assembly and motility, determining the underlying basis for disease can be challenging. The study of model organisms with motile cilia like *Chlamydomonas* and *Tetrahymena* has proven useful for understanding the network of genes and proteins required for motility.

Comparative genomics and proteomics have identified >1000 proteins as structural components of the ciliary axoneme and ciliary membrane (Sakoto-Antoku and King, 2022, reviewed in van Dam et al., 2019). Most motile cilia contain nine outer doublet microtubules (DMTs) surrounding two singlet central pair (CP) MTs. Each DMT contains thousands of large, multisubunit motors known as the outer and inner dynein arms (ODA, IDA). The dynein motors generate the force for microtubule sliding between the A-tubule of one DMT and the B-tubule of the adjacent DMT. The ODAs and IDAs are composed of different heavy, intermediate, and light chain (DHC, IC, LC) subunits attached in two rows on the A-tubule. They are further organized into functional units that repeat every 96 nm, with four identical ODAs and seven unique IDAs (I1/*f, a, b, c, e, g, d*) bound at discrete sites within the repeat. Motor activity is coordinated in part by signals from the CP and its numerous projections that contact three radial spokes (RS1, RS2, RS3 or RS3S) bound near the base of the IDAs in each 96 nm repeat (reviewed in Brown et al., 2023; Witman and Mitchell, 2023). The IC/LC complex of the I1/f dynein forms a regulatory node at the base of RS1, the nexin-dynein regulatory complex (N-DRC) at the base of the RS2, and a calmodulin spoke complex (CSC) connects between the bases of RS2, RS3/RS3S and the N-DRC (Heuser et al., 2009; 2012a, b). The I1 dynein and N-DRC also connect to the ODAs and other structures in the 96 nm repeat to stabilize the binding of the IDAs and coordinate dynein activity (Heuser et al., 2009; 2012; Oda et al., 2013). This network includes the tether/tether head (T/TH) structures associated with the I1 dynein motor domains (Fu et al., 2018; Kubo et al., 2018), the MIA complex between the I1 dynein and the ODAs (King and Dutcher, 1997; Yamamoto et al., 2013), and the FAP57 complex linking the MIA complex, IDAs *g* and *d*, and the T/TH structure (King et al., 1994; Lin et al., 2019; Bustamante-Marin et al., 2020; Ghanaeian et al., 2023).

Many mutations that disrupt the assembly of ODAs, IDAs, RSs, CP and its projections, MIA complex, N-DRC, T/TH, and FAP57 complex have been characterized over decades of work in *Chlamydomonas*, *Tetrahymena*, and other organisms, and their motility phenotypes are very diverse, ranging from flagellar paralysis, ciliary twitching, reduced beat frequency, and/or altered ciliary waveforms with reduced amplitudes (reviewed in Sale and Dutcher, 2023). However, ciliary motility is extremely complex, and many organisms can adjust the pattern of their waveforms in response to internal and external stimuli. For instance, *Chlamydomonas* can adjust the beat frequency of its *cis* and *trans* cilia to undergo both positive and negative phototaxis, or it can reverse its swimming direction in response to bright light (reviewed in Witman, 1993; Dutcher, 2020; Sale and Dutcher, 2023). The latter involves the conversion of an asymmetric ciliary-type waveform to a more symmetric, flagellar-type waveform. The structures and polypeptides responsible for waveform conversion are poorly understood. A handful of mutations result in cells that swim forward with a symmetric waveform (*pf10, pf12/pacrg*, and *fap20*) (McVittie, 1972; Dutcher et al., 1988; Frey et al., 1997; Dymek et al., 2019), and a second group result in cells that swim backwards with a symmetric waveform (*mbo1, mbo2*, and *mbo3*) (Segal et al., 1984). Characterization of the *fap20* and *pacrg/pf12* mutants by conventional transmission electron microscopy (TEM) and cryo-electron tomography (cryo-ET) has shown that FAP20 and PACRG form the inner junction between the A- and B-tubules of the DMT (Yanagisawa et al., 2014; Dymek et al., 2019). These studies also revealed secondary defects in the assembly of structures located inside the lumen of the proximal B-tubule (beak-MIP), the IDA *b*, and the I1 dynein T/TH complex (Yanagisawa et al., 2014; Dymek et al., 2019). Previous studies of *mbo* mutants described defects in the assembly of eight axonemal polypeptides ranging in size from ∼33-245 kD, but the identities and locations of the missing proteins were unknown (Segal et al., 1984). The only structural defect identified by conventional TEM was the absence of beak structures located within the lumens of the B-tubules of DMT5 and DMT6. The isolation and characterization of new alleles of *MBO2* by insertional mutagenesis later revealed MBO2 to be a conserved, coiled coil protein found along the length of the axoneme (Tam and Lefebvre, 1993, 2002).

Here, we re-analyzed the *mbo* mutants to identify the polypeptides and structures responsible for waveform conversion and backwards swimming. We used proteomic approaches and tagged *MBO2* transgenes to identify a large complex of polypeptides missing or reduced in the *mbo* mutants and restored in *MBO2* rescued strains. Comparative proteomics revealed that several of these proteins are also altered in *pf12/pacrg* and *ida8*/*fap57* mutants. Cryo-ET revealed the complexity of doublet-specific structural defects, including an elongated, L-shaped structure reaching between the base of IDA *b* and other axonemal complexes. Rescue with SNAP-tagged MBO2 constructs followed by streptavidin gold labeling and classification averaging suggests that MBO2 is one of the components of the L-shaped structure that interconnects multiple regulatory components and targets the assembly of IDA *b* within the 96 nm axoneme repeat.

## Results

### Identification of polypeptide defects in mbo2 and mbo1 axonemes

Transformation of *mbo2* with an HA-tagged *MBO2* transgene rescued the *mbo* motility phenotype, increased forward swimming velocities, and restored assembly of the missing MBO2 subunit along the length of the axoneme (Figure 1A-C, Supplementary Figure S1, Supplementary Tables S2, S3, see also Tam and Lefebvre, 2002). To identify polypeptides associated with MBO2, we purified axonemes from wild-type, *mbo2*, *MBO2-HA* rescued strains, and *mbo1,* labeled them using iTRAQ isobaric tags, and subjected them to quantitative mass spectrometry (see Materials and Methods for details). Nineteen proteins were reduced below 75% (P value < 0.05) in four independent iTRAQ experiments using *mbo2* and *mbo1* (Supplementary Table S4). We later modified the protocol using three independent samples of WT, *mbo2*, and *MBO2-HA* axonemes, TMT-based isobaric tags, and more sensitive instrumentation (Supplemental Figure S2A, Supplemental Table S4). All 19 proteins identified above were reduced in *mbo2* and restored in *MBO2-HA* in the TMT experiment, but only 10 proteins were significant at P values < 0.05 (Table 1). Several very low abundance proteins not previously detected by the iTRAQ experiments were also identified as significantly reduced in *mbo2* and restored in MBO2-HA in the TMT experiment (Supplemental Table S4).

**Figure 1.**
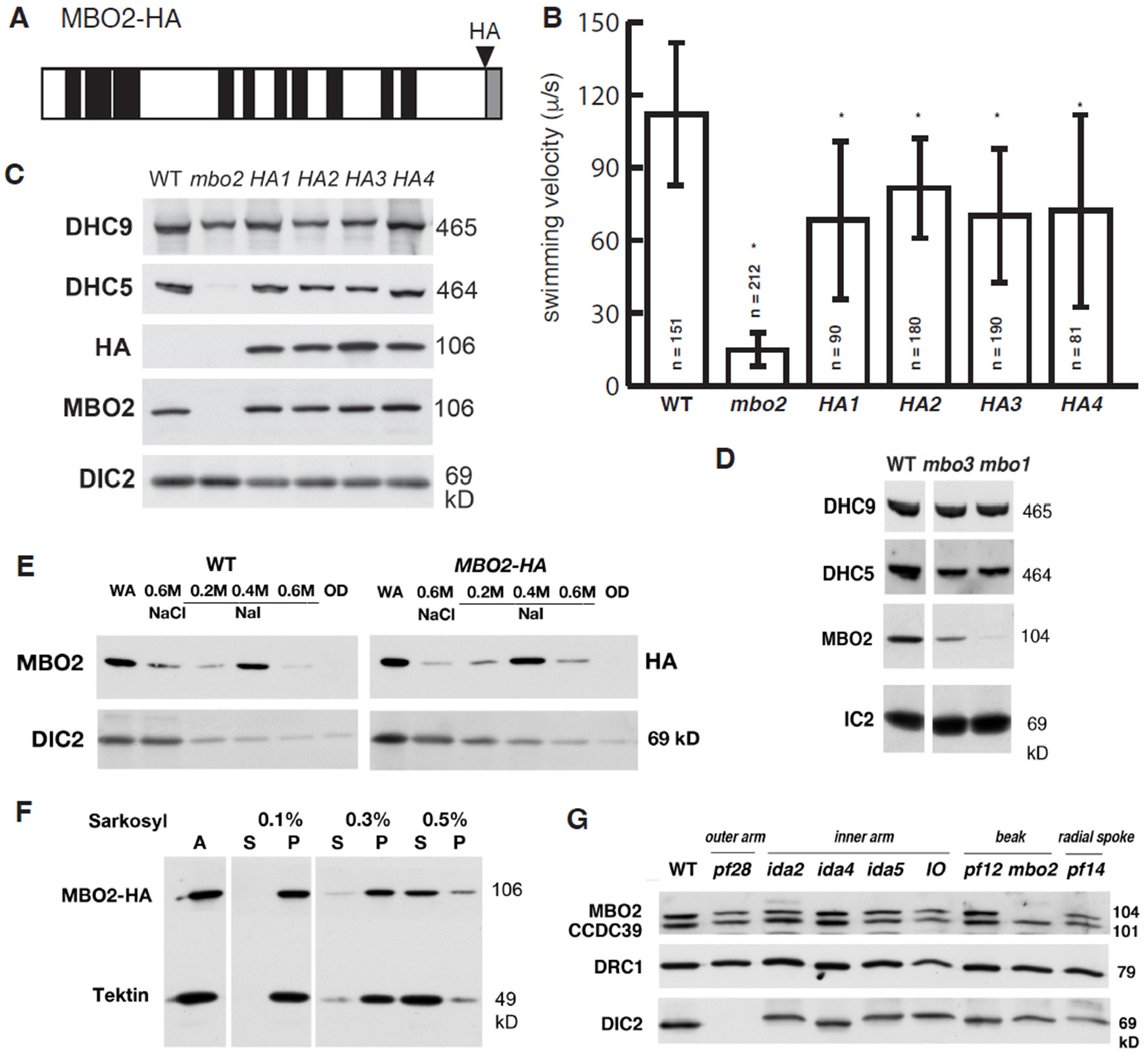
MBO2 forms a complex that stabilizes assembly of DHC5. **(A)** A schematic diagram of the MBO2 polypeptide showing the location of coiled coil domains (black) and a disordered region in the C-terminus gray). Also shown is the position of the HA epitope tag. (**B**) Transformation of *mbo*2 with *MBO2-HA* restores forward swimming. The forward swimming velocities of WT, *mbo2*, and *HA*-rescued strains (*HA1-HA4*) were measured by phase contrast light microscopy. All the *HA*-rescued strains swam forwards significantly faster (P<0.5) than *mbo2*, but significantly slower than WT, consistent with previous reports (Tam and Lefebvre, 2002). (**C**) Western blots of axonemes probed with different antibodies show the recovery of both MBO2 and DHC5 in the *HA*-rescued strains. DHC9 and DIC2 are loading controls for other IDA and ODA isoforms. (**D**) Western blots of *mbo1* and *mbo3* axonemes show that MBO2 and DHC5 are also reduced in these two strains. (**E**) WT and *mbo2; MBO2-HA* axonemes were subjected to sequential extraction with 0.6M NaCl and 0.2-0.6M NaI buffers, and analyzed on Western blots (WA, whole axoneme; OD, extracted outer doublets). The majority of MBO2 was extracted with 0.4M NaI, whereas dynein subunits (DIC2) were largely extracted with 0.6M NaCl. **(F)** Western blot of WT axonemes (A) that were extracted with increasing concentrations of Sarkosyl (S, supernatant; P, pellet). (**G**) Western blot of axonemes from different motility mutants lacking outer arms (*pf28*), inner arms (*ida2, ida4, ida5*), B-tubule beaks (*pf12, mbo2*) and radial spokes (*pf14*).

**Table 1.** Protein ratios in mbo2 and MBO2-HA axonemes determined by TMT labeling and MS/MS.

| Protein (Cre) | Peptides | *mbo2*/WT | HA/WT |
| --- | --- | --- | --- |
| **MBO2**<br>Cre09.g416550 | 55 | **0.12** | 1.00 |
| **FAP58\***<br>Cre13.g584300 | 65 | **0.12** | 1.04 |
| **FAP324**<br>Cre08.g383101 | 11 | **0.08** | 0.93 |
| **FAP51**<br>Cre02.g077600 | 81 | **0.21** | 1.08 |
| **FAP331**<br>Cre06.g308000<br>FAP57 paralog | 48 | **0.18** | **0.54** |
| **DHC5**<br>Cre02.g107050 | 120 | **0.36** | 0.92 |
| **Cre16.g682850** | 30 | **0.30** | 1.14 |
| **FAP238**<br>Cre12.g509350 | 17 | **0.38** | 1.15 |
| **FAP271**<br>Cre12.g486700 | 17 | **0.40** | 1.00 |
| **FAP308**<br>Cre10.g430700 | 65 | **0.40** | 0.97 |
| **Cre07.g313850**<br>FAP337 paralog | 31 | 0.50 | 0.73 |
| **FAP343**<br>Cre06.g278234 | 29 | 0.52 | 1.01 |
| **FAP258**<br>Cre07.g325751 | 32 | 0.52 | 1.01 |
| **FAP395**<br>Cre06.g275950 | 47 | 0.43 | 1.04 |
| **FAP327**<br>Cre01.g040600 | 39 | 0.43 | 1.06 |
| **CPY**<br>Cre12.g483800 | 35 | 0.51 | 1.01 |
| **Cre05.g234640** | 45 | 0.50 | 1.11 |
| **Cre07.g354650** | 53 | 0.63 | 0.94 |
| **Cre01.g029800** | 29 | 0.59 | 0.93 |
| **Other DMT proteins** |  |  |  |
| DRC4 | 54 | 0.95 | 1.06 |
| PACRG | 23 | 1.04 | 0.96 |
| FAP59 | 67 | 1.21 | 1.09 |
| FAP172 | 54 | 1.16 | 1.07 |
| FAP57 | 62 | 1.36 | 1.20 |
| FBB7 | 47 | 0.69 | 0.89 |
| FAP337 | 25 | 0.45 | 1.16 |
| FAP73 | 18 | 1.10 | 1.03 |
| FAP100 | 26 | 1.11 | 1.02 |
| FAP78 | 62 | 1.11 | 1.06 |
| FAP189 | 76 | 1.22 | 0.99 |
| RSP16 | 24 | 1.03 | 0.99 |
| RIB43 | 35 | 1.13 | 1.03 |
| RIB72 | 39 | 1.09 | 0.97 |
| TUA | 24 | 1.01 | 0.98 |
| TUB | 25 | 1.00 | 1.00 |
| Tektin | 25 | 1.55 | 1.36 |

One unexpected finding was a ∼60-70% reduction of DHC5 in *mbo2* axonemes. DHC5 is one of twelve DHCs forming the inner dynein arms (IDA) in *Chlamydomonas* (reviewed in King et al., 2023). To confirm the decrease of DHC5, we probed Western blots of WT and mutant axonemes with DHC5 and DHC9 specific antibodies. DHC5 was reduced in *mbo2* and restored to WT levels in *MBO2-HA* rescued axonemes, whereas DHC9 was present at WT levels in all samples (Figure 1C). Proteomic analysis of other inner arm DHCs indicated no significant changes in the DHC ratios between WT, *mbo2,* and *MBO2-HA* axonemes (Supplemental Table S4). DHC5 was also reduced ∼25-40% in *mbo1* and *mbo3* axonemes (Figure 1D, Supplemental Table S4), which suggests that loss of MBO2 destabilizes the attachment of DHC5 to the DMT. To better understand the relationship between DHC5 and MBO2, we tested whether these polypeptides might co-fractionate following biochemical extraction of purified axonemes. The IDAs can be extracted from WT axonemes by treatment with 0.6M NaCl or KCl (Pazour et al., 1995), but extraction of MBO2 required treatment with 0.4-0.6M NaI or 0.5% Sarkosyl (Figure 1E, F), indicating that they are in distinct biochemical complexes. A subset of the proteins reduced in *mbo2* are enriched in 0.6M NaCl/0.6M KCl extracts (Table 2, Supplemental Table S4, Pazour et al., 2005), suggesting that they might be associated with DHC5. The 0.6M NaCl extracts can be partially purified by ion exchange FPLC based chromatography into seven distinct peaks containing different IDA isoforms (Kagami and Kamiya, 1992). Proteomic analysis of the DHC5-containing fraction identified several polypeptides, but only one of these proteins, Cre14.g618300, was identified as reduced in *mbo2* by proteomics (Supplemental Figure S2B, Supplemental Table S4). These results indicate that most MBO2-associated proteins are dissociated from DHC5 during high salt extraction and FPLC fractionation.

**Table 2.** Characteristics of proteins reduced in *mbo2* and *mbo1* axonemes.

| Protein Name<br>(JGI Gene ID<br>Phytozome13) | AA<br>(MW) | Predicted<br>Domains | *Volvox*<br>*carteri*<br>JGI ID<br>(Blastp) | *Physcomitrella*<br>*patens*<br>Genbank ID<br>(Blastp) | *Homo sapiens*<br>Genbank ID<br>(Blastp) |
| --- | --- | --- | --- | --- | --- |
| **MBO2 core subunits (only reduced in *mbo*, abundant, enriched in extracted axonemes)** |  |  |  |  |  |
| **MBO2**<br>Cre09.g416550 | **920**<br>(104 kD) | CC, low<br>complexity | 0011s0271<br>0.0 | PNR44978.1<br>(1e-100) | **CCDC146**<br>NP_065930.2<br>(1e-102) |
| **FAP58\***<br>Cre13.g801543<br>Cre13.g584300 | **871**<br>(102 kD) | CC, low<br>complexity | 0001s0063<br>0.0 | **CFAP58L**<br>XP_024400258.1<br>(0.00) | **CCDC147**<br>NP_0010088723.1<br>(0.0) |
| **FAP51**<br>Cre02.g077600 | **1939**<br>(208 kD) | CC | 0030s0082<br>(0.0) | XP_023438985.1<br>(1.9) | ND |
| **DHC5 pattern (also reduced in short or *pf12/pacrg* axonemes, enriched in high salt extracts)** |  |  |  |  |  |
| **DHC5**<br>Cre02.g107050 | **4185**<br>(464 kD) | AAA, CC | 0022s0148<br>(0.0) | **DHC7**<br>XP_024376875.1<br>(0.0) | **DNAH7**<br>AAL37427.1<br>(0.0) |
| **FAP343**<br>Cre06.g278234 | **1079**<br>(114 kD) | CC, WW | 0002s0194<br>(2.2e-140) | **CEP164L**<br>XP_024400159.1<br>(7e-12) | **CEP164**<br>BAG64184.1<br>(5e-10) |
| **FAP324**<br>Cre08.g383101 | **631**<br>(70 kD) | CC, Kelch,<br>FN3 | 0011s0100<br>(0.0) | XP-024360494.1<br>(2e-22) | **KLHDC3**<br>EAX04137.1<br>(1e-12) |
| **FAP238**<br>Cre12.g509350 | **366**<br>(42 kD) | EF-hand,<br>CC | 0016s0228<br>(3.3e-154) | XP_02439029.1<br>(0.095) | CAA27243.1<br>(0.003) |
| ***pf12* pattern (also reduced in *pf12* axonemes, not enriched in high salt extracts or unknown)** |  |  |  |  |  |
| **Cre07.g354650** | **3624**<br>(357 kD) | CC, cNMP | 0005s0121<br>1.6e-100 | ND | ND |
| **Cre16.g682850** | **864**<br>(87 kD) | CC, low<br>complexity | 0018s0102<br>1e-68 | ND | ND |
| **FAP258**<br>Cre07.g325751 | **754**<br>(74 kD) | CC | 0003s0163<br>(4.8e-74) | ND | ND |
| **FAP271**<br>Cre12.g486700 | **384**<br>(40 kD) | Low<br>complexity | 0016s0163<br>(4.3e-141) | **STPG2L**<br>XP_024367883.1<br>(0.052) | **STPG1**<br>EAW95125.1<br>(2e-4) |
| **FAP57 complex (altered in *mbo* and *fap57* (*ida8*) axonemes, enriched in high salt extracts)** |  |  |  |  |  |
| **FAP331**<br>Cre06.g308000<br>FAP57 paralog | **1989**<br>(205 kD) | WD, CC | 0009s0155<br>(0.0) | **CFAP57L**<br>XP_024388298.1<br>(4e-07) | **CFAP57**<br>(3e-13) |
| **Cre07.g313850**<br>FAP337<br>paralog | **1309**<br>(136 kD) | WD, EF<br>hand | 0003s0128<br>(0.00) | **WDY (WDR49)**<br>XP_024383320.1<br>(2e-79) | **WDR49**<br>NP_001335880.1<br>(6e-64) |
| **Enriched in short axonemes, poorly conserved, variable abundance** |  |  |  |  |  |
| **FAP395**<br>Cre06.g275950 | **2136**<br>(219 kD) | CC | 0009s0155<br>(3.2e-109) | XP_024367482.1<br>(5e-06) | ND |
| **FAP327**<br>Cre01.g040600 | **1794**<br>(186 kD) | CC | 0001s1006<br>(2.2e-154) | XP_024363033.1<br>(0.022) | ND |
| **(CPY)**<br>Cre12.g483800 | **2528**<br>(259 kD) | CC, low<br>complexity | 0031s0129<br>(1.5e-55) | XP_024392807.1<br>(3.9) | ND |
| **Cre05.g234640** | **3529**<br>(348 kD) | IQ, low<br>complexity | 008s0412<br>(4.2e-51) | ND | ND |
| **Cre01.g029800** | **963**<br>(101 kD) | CC | 0001s0930<br>(4.3e-172) | ND | ND |
| **FAP308**<br>Cre10.g430700 | **3085**<br>(315 kD) | CC | 0005s0270<br>(9e-118) | ND | ND |

### Characterization of other mutants that share overlapping polypeptide defects with mbo strains

Because IDA *b* is reduced in *pacrg* and *pacrg; fap20* axonemes (Dymek et al, 2019), we analyzed the proteome of a *pacrg* mutant, *pf12*, for potential overlap with proteins missing in *mbo1* and *mbo2*. The *pf12* mutation disrupts assembly of PACRG, one of two proteins that form the inner junction between the A- and B-tubules of the DMT (Dymek et al., 2019). The *pf12/pacrg* strains swim forwards with a symmetric waveform, lack beaks inside the B-tubules of DMTs 5 and 6, and assemble reduced levels of IDA *b* into the axoneme (McVittie, 1972; Frey et al., 1997; Dymek et al., 2019). Proteomics of *pf12* axonemes confirmed that PACRG is reduced to <10% of WT levels, whereas MBO2 is essentially WT level, as also shown by Western blot (Supplemental Table S4, Figure 1G) (Dymek et al., 2019). However, nine MBO2-associated proteins were significantly and consistently reduced to <75% of WT levels in *pf12* axonemes (Supplemental Table S4, Table 2). These proteins may include potential docking factors for DHC5 or subunits of other complexes destabilized in both *mbo* and *parcg* mutants.

FAP331 and Cre07.g313850 are two proteins reduced in *mbo1, mbo2,* and *pf12* that are paralogs of related proteins in the FAP57 complex and that are increased in *ida8/bop2/fap57* axonemes (Lin et al., 2019; Bustamante-Marin et al., 2020). We analyzed the axonemes from *mbo1, mbo2, pf12* and a *fap57* mutant to clarify potential interactions between the different protein complexes (Supplemental Table S4). The *ida8-1* mutation disrupts assembly of FAP57, and FAP337 and three IDAs were also reduced in *ida8* axonemes (Lin et al., 2019). These defects were partially offset by increases in the levels of two minor paralogs of FAP57, FBB7 and FAP331, and one paralog of FAP337, Cre07.g313830, in *ida8/bop2/fap57* (Lin et al., 2019; Bustamante-Marin et al., 2020; and Supplemental Table S4). No other MBO2-associated proteins were altered in *ida8/fap57* axonemes (Supplemental Table S4), and FAP57 ratios were WT in both *mbo* and *pf12/pacrg* mutants (Table S4). However, FBB7 was reduced in *pf12*, and FAP337 was reduced in *pf12* and *mbo2,* but not *mbo1*. (Supplemental Table S4, Table 2). These results suggested that changes in the minor FAP57 paralogs and associated FAP337/Cre07.g313850 subunits might be secondary defects in *mbo* and *pf12* mutants.

We also analyzed the proteome of a *pf9-2; pf28* double mutant strain with short axonemes (∼2.9 μm) to identify polypeptides that might be enriched in the proximal or distal regions of the axoneme (Hwang et al., submitted). DHC5 is reduced in the proximal 2 μm of the axoneme and reduced in short axonemes during flagellar regeneration (Yagi et al., 2009). Inspection of the short axoneme proteome confirmed that the proximal DHCs (DHC3, DHC4, DHC11) were increased, and DHC5 was reduced (Hwang et al., submitted; Supplemental Table S4). Three other MBO2-associated proteins and FAP331 were reduced in short axonemes (<60%), whereas four MBO2-associated proteins were increased (>140%) (Supplemental Table 4). These results suggest that some of MBO2-associated proteins may vary between the proximal and distal axoneme.

### Phenotypic characterization of other mutants with defects in DHC5/IDA b

Given the decrease in DHC5 observed in *mbo* axonemes, we analyzed other motility mutants with DHC5 defects. The *IO* mutant (CC-2012) is an isolate of *pf23* that assembles full-length, paralyzed flagella lacking most of the IDAs, including IDA *b*/DHC5. However, Western blots showed that MBO2 levels were WT (Figure 1G). We also searched the CLiP library of insertional mutants (Li et al., 2016; 2019) to identify potential *dhc5* mutations (Supplemental Table S1). PCR of genomic DNA identified four strains with plasmid insertions into the 5’ region of the *DHC5* gene (Figure S3A-C). Western blots of axonemes probed with a DHC5 antibody detected bands migrating at the sizes predicted for N-terminal fragments containing the tail domain but lacking the C-terminal motor domain. MBO2 was also present at WT levels (Supplemental Figure S3D). Analysis of swimming phenotypes showed that the *dhc5* strains swam forwards at WT speeds with no obvious motility defects (Supplemental Figure S3E), demonstrating that absence of the DHC5 motor domain/activity itself does not result in a *mbo* phenotype.

### Cryo-electron tomography reveals new structural defects in mbo2 axonemes

Studies of *mbo* axonemes by conventional TEM have shown defects in the assembly of beak structures within the lumens of the B-tubules of DMT5 and DMT6 but failed to reveal any defects in the assembly of the IDAs (Segal et al., 1984; Tam and Lefebvre, 2002). We used cryo-ET to re-analyze the axonemal structure, including the IDAs, in WT and *mbo2* axonemes (Figure 2). Subtomogram averages of 96 nm repeats from all WT DMTs revealed the two-headed I1/*f* dynein at the proximal end of the repeat, followed by the six, single-headed IDAs (*a, b, c, e, g, d*) (Figure 2A-D). However, the electron density of IDA *b*, which contains DHC5 and locates just distal to RS1, was weaker than for other IDAs, suggesting IDA *b* was not present in all WT 96 nm repeats (Figure 2A-D). In contrast to WT, tomograms of the 96nm repeat from all DMTs in *mbo2* lack IDA *b* almost entirely (Figure 2E-H). As previously reported for WT axonemes, IDA *b* is significantly reduced on all DMTs in the proximal region of the axoneme (Yagi et al., 2009) and specifically reduced on certain DMTs in the medial/distal region (Bui et al., 2012; Lin et al., 2012, 2019; Dymek et al., 2019). Therefore, we performed classification analyses to determine the presence or absence of IDA *b* in different regions of the axoneme and on specific DMTs (Figure 2I, J). In WT, IDA *b* was present in 6% of the proximal tomograms and 55% of the medial/distal tomograms; in *mbo2*, IDA *b* was only present in 2% of the proximal tomograms and 12% of the medial/distal tomograms (Figure 2I). In WT axonemes, IDA *b* was reduced on all DMTs in the proximal region and on DMT1, DMT5, and DMT9 in the medial/distal region (Figure 2J), consistent with previous reports (Bui et al., 2012, Lin et al., 2012, 2019; Dymek et al., 2019). In *mbo2*, IDA *b* was reduced in all DMTs, with the most significant changes from WT in the medial/distal region of DMT2 to DMT8 (Figure 2J). Examples of the distribution of IDA *b* in individual axonemes are shown in Figure 2K.

**Figure 2.**
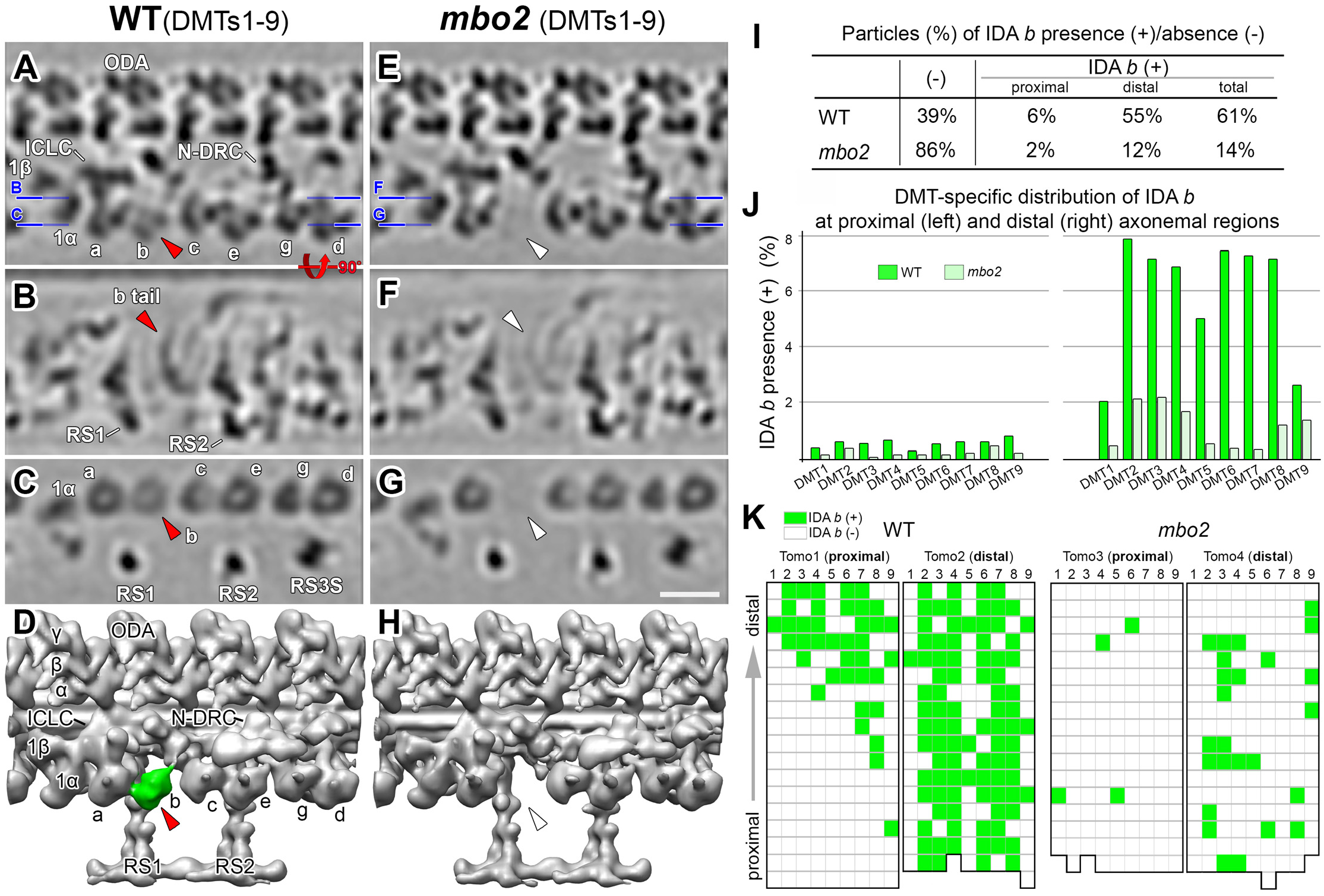
IDA *b* is reduced in the 96 nm axonemal repeat from *mbo2*. **(A–H)** Tomographic slices (A–C, E–G) and isosurface renderings (D, H) of the averaged 96-nm repeats from all DMTs of WT (A–D) and *mbo2* (E–H) axonemes. Blue lines in (A and E) indicate the locations of the slices in the respective panels. IDA *b* can be visualized in WT (red arrowheads) but not in *mbo2* (white arrowheads). **(I)** The presence (+) and absence (-) of IDA *b* in the 96nm repeats from the proximal and medial-distal regions of WT and *mbo2* axonemes. **(J)** The bar graph shows the fraction of repeats with IDA *b* for each DMT from WT (green) and *mbo2* (light green) in the proximal and distal regions. **(K)** Distribution pattern of IDA *b* in four tomograms of proximal and distal regions from WT (left) and *mbo2* (right). Each grid represents a single repeat, and the colors indicate whether IDA *b* is present (green) or absent (white) in the repeat. From bottom to top, the grids represent the repeats in the proximal-to-distal direction (gray arrow). Note that IDA *b* is not present in the proximal region and Tomo 1 of WT shows a transition from the proximal-to-distal region. Other labels: 1α, β, the I1 dynein α- and β-head; a-g, single-headed IDAs; ICLC, intermediate chain and light chain complex of I1 dynein; IDA, inner dynein arm; N-DRC, nexin-dynein regulatory complex; ODA, outer dynein arm; RS1/2/3S, radial spokes 1, 2 and 3 stand-in. Scale bar in G, 20 nm (valid for A–C and E–G).

Because several polypeptides are missing or reduced in *mbo2* axonemes, we analyzed the region around IDA *b* using DMT specific averaging (Figure 3). The averages of DMT1 and DMT9 did not show significant changes between WT and *mbo2* (Figure 3C, K). These results are consistent with the observation that IDA *b* was rarely found on these DMTs in either WT or *mbo2* (Figure 2J). For DMT5 to DMT8, both IDA *b* and the WT structures near the base of IDA *b* were completely missing in *mbo2* (Figure 3G-J). This suggests that a subset of the proteins missing in *mbo2* are located within the structures around the base of IDA *b*. In contrast, the *mbo2* defects were different in the averages of DMT2 to DMT4. Even though IDA *b* was missing in *mbo2*, the region surrounding the base of IDA *b* retained significant structural density, similar but not identical to the structures seen in WT (compare purple/blue density in Figure 3D-F). These structures may contain other proteins that are related to proteins missing in *mbo2*, but do not depend on MBO2 for assembly into the axoneme.

**Figure 3.**
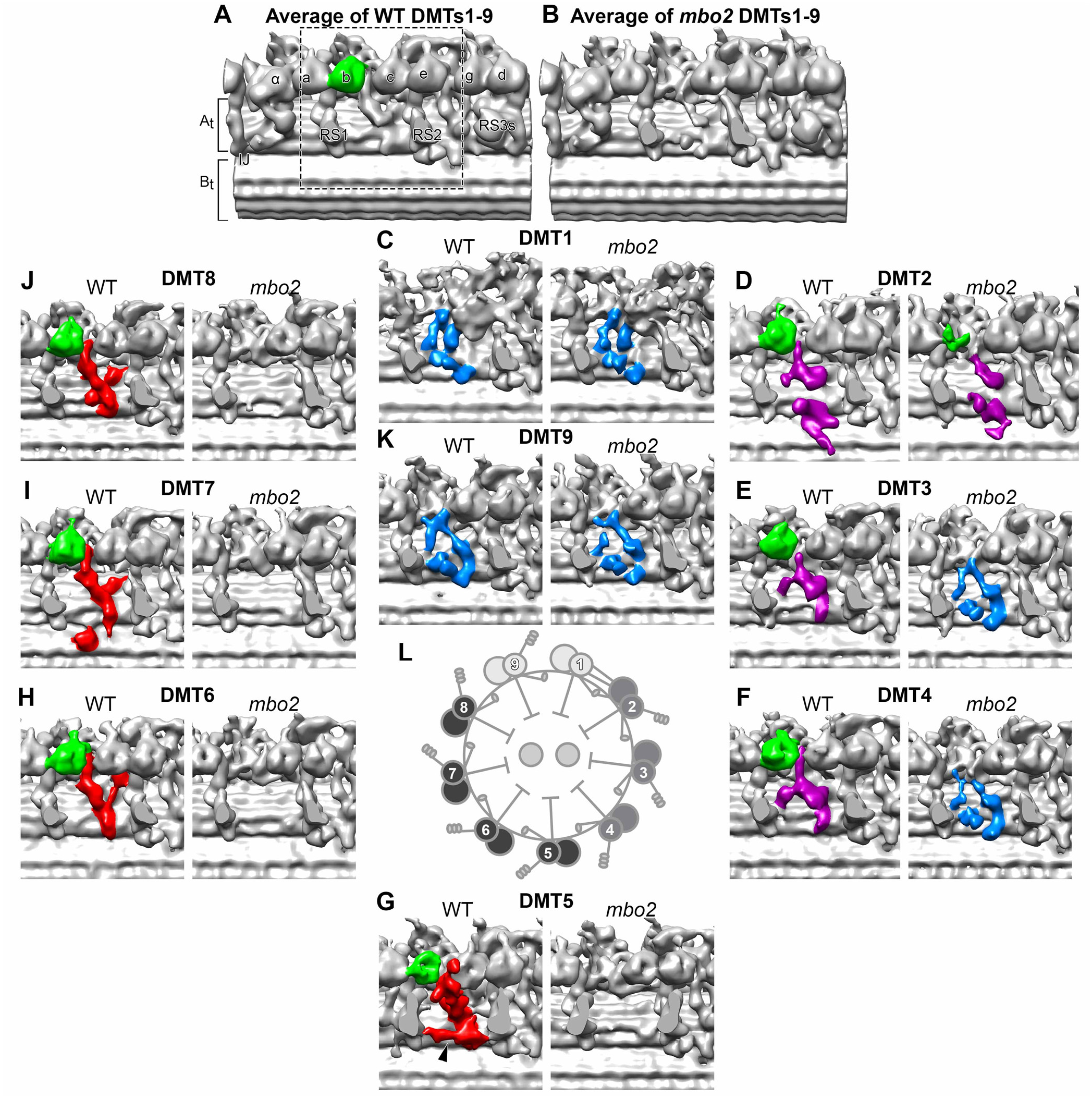
Doublet microtubule (DMT) specific defects in *mbo2*. **(A, B)** Bottom view of the 3D isosurface renderings of the averaged 96-nm repeats from all DMTs in WT (A) and *mbo2* (B) axonemes. Radial spokes (RS) were cropped to allow an-unhindered view of the IDA-to-DMT-docking region. Motor domain of IDA b is colored green **(C–K)** Comparison of the IDA *b* tail-associated structures across nine DMTs between WT (left) and *mbo2* (right) focusing on the region marked by the dashed box in A. Based on the structural characterizations in WT and defects in *mbo2*, the WT DMTs were categorized into three groups: DMTs1, 9 (blue), DMTs2-4 (purple) and DMTs5-8 (red). The black arrowhead in WT DMT5 indicates an DMT5-specific structure connecting RS1 and RS2. **(L)** A schematic drawing of the axoneme in cross-section to show the asymmetric distribution of the IDA *b* associated structures on the three classes of DMTs.

Tomograms from the proximal region of the axoneme were also analyzed for the presence or absence of the beak structures inside the B-tubules of DMT1, DMT5, and DMT6 (Supplemental Figure S4). Beak structures were visible in DMT1 and DMT6 from both WT and *mbo2* axonemes (Figure S4A-B, E-F) but not visible in DMT5 from *mbo2* axonemes (Supplemental Figure S4C-D).

### Rescue of mbo2 defects and mapping the position of MBO2 by transformation with SNAP-tagged MBO2 transgenes

The large number of MBO2-associated proteins and the complexity of structural defects in *mbo2* axonemes make precise localization of the MBO2 subunit impossible using a WT-mutant comparison. Therefore, we constructed three *MBO2* transgenes with SNAP tags located at the N-terminus (N-SNAP), near the middle of the protein (M-SNAP at amino acid 569) and at the C-terminus (C-SNAP) (Figure 4A, Supplemental Figure S1). Each transgene was introduced into *mbo2* by co-transformation, and transformants were screened for recovery of forward swimming and re-assembly of MBO2 and DHC5 into the axoneme. Western blots of axonemes from rescued strains demonstrated that MBO2 was assembled at WT levels and migrated at the size expected for a SNAP-tagged subunit. Assembly of DHC5 was also restored in the rescued strains (Figure 4B). Measurement of swimming velocities indicated the N-SNAP and C-SNAP rescued strains recovered ∼72% of WT velocity, whereas the M-SNAP rescued strain recovered only ∼48% of the WT speed (Figure 4C). However, analysis of cell trajectories and motility by phase contrast microscopy confirmed that all the rescued strains were swimming forwards with asymmetric waveforms (Figure 4D). The SNAP-tagged transgenes therefore restored sufficient function to rescue the *mbo* phenotype, even though the rescued cells were not completely WT with respect to swimming velocities.

**Figure 4.**
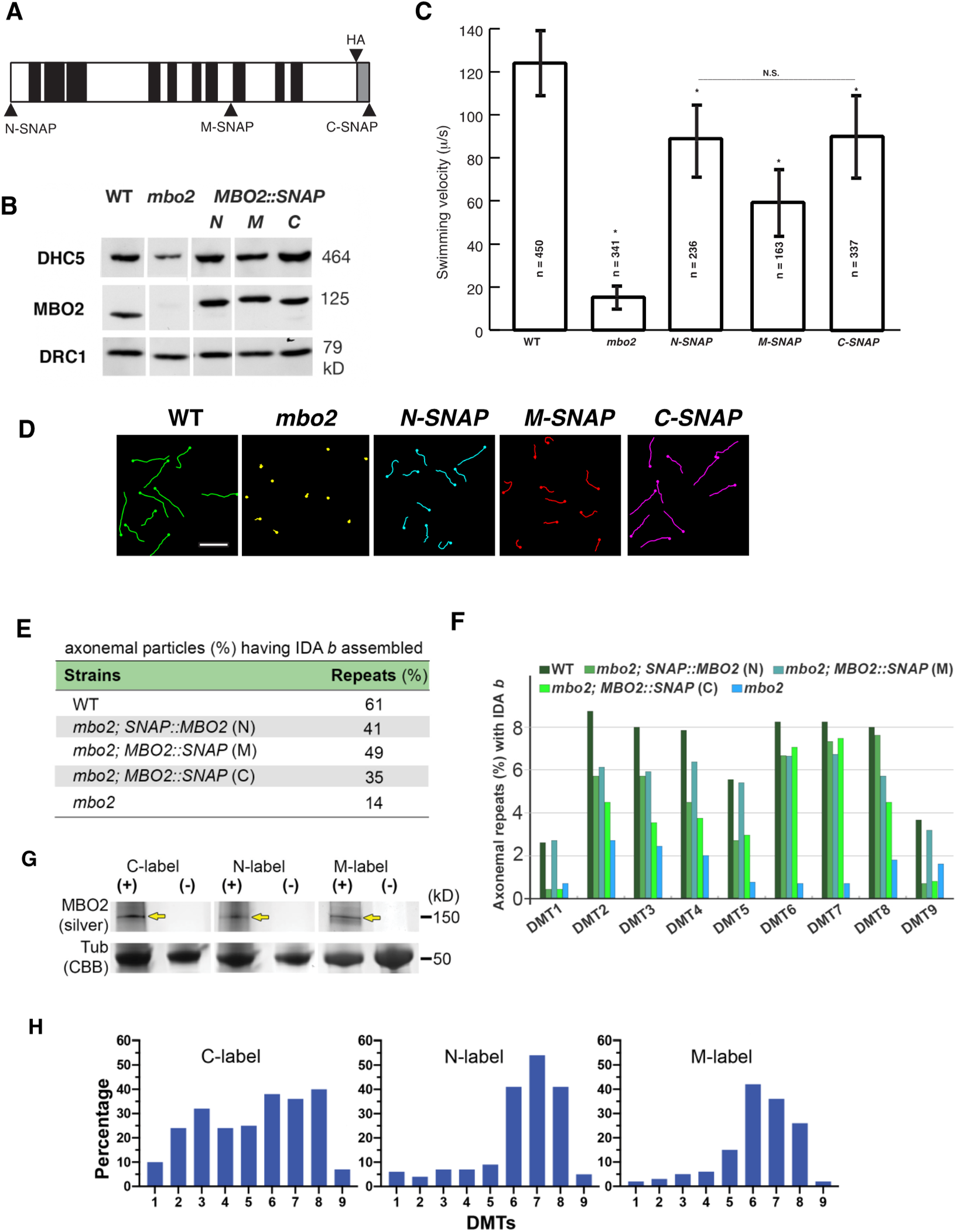
Rescue of biochemical, motility, and structural defects in the SNAP-tagged *MBO2* strains. (**A**) Schematic diagram of MBO2 showing the insertion of SNAP tags at the N-terminus (N-SNAP, M1), in the middle (M-SNAP, L569) and at the C-terminus (C-SNAP, I920) of the polypeptide. Also shown are the coiled coil domains (black), disordered region (gray) and HA tag. (**B**) Western blot of axonemes from WT, *mbo2*, and *MBO2::SNAP* rescued strains (N, M, C) probed with different antibodies. Both DHC5 and MBO2 are re-assembled in the rescued strains; DRC1 is a loading control. (**C**) Transformation of *mbo2* with each construct significantly increased (P<0.05) swimming velocity relative to *mbo2* (see asterisks). *N-SNAP* and *C-SNAP* were significantly faster than *M-SNAP*, slower than WT, but not significantly different from one another. (**D**) Forward swimming trajectories of each strain are shown here for an interval of 1 second. (The *mbo2* strain is swimming backwards.) Scale bar, 50 μm. **(E)** Percentage of 96 nm axoneme repeats that contain IDA *b* in WT, *mbo2*, *mbo2; MBO2::SNAP* (N), *mbo2; SNAP::MBO2* (M) and *mbo2; MBO2::SNAP* (C) strains. The assembly of IDA *b* is increased in the rescued strains as compared to *mbo2*. **(F)** Bar graph shows the percentage of 96 nm repeats that contain IDA *b* for each DMT from WT, the three SNAP-tagged rescued strains, and *mb*o*2*. IDA *b* is increased on DMTs 2-8 in all rescued strains as compared to *mbo2*. **(G)** SDS-PAGE of SNAP-tagged axonemes labeled with streptavidin gold and detected by silver enhancement. The yellow arrows mark the SNAP-tagged MBO2 polypeptides. Tubulin was stained with Coomassie Brilliant Blue (CBB) as a loading control. **(H)** Percentage of 96 nm repeats with gold-labeled SNAP-tags found on each DMT in *mbo2; MBO2::SNAP* (C), *mbo2; SNAP::MBO2* (N) and *mbo2; MBO2::SNAP* (M).

To obtain a more quantitative measurement of recovery, we analyzed the DHC composition of the rescued strains by SDS-PAGE, mass spectrometry, and cryo-ET. Axonemes from WT, *mbo2-4*, and the three SNAP-tagged rescued strains were fractionated by SDS-PAGE, and the DHC region was excised for MS/MS analysis (see boxed region in Supplemental Figure S2C). The relative abundance of each IDA DHC was estimated by spectral counting and plotted as a percentage of DHC content of the I1 dynein (Supplemental Figure S2D). Quantification of peptides using the tools available in Proteome Discover yielded similar results (Supplemental Table S4). Only DHC5 was significantly reduced in *mbo2*, and it was re-assembled in the SNAP-tagged rescued axonemes at 64-84% of WT levels. Inspection of tomograms from the medial and distal regions of axonemes confirmed that IDA *b* was present in 14% of the 96 nm repeats in *mbo2* and increased to 35-49% of the repeats in the SNAP-tagged rescued strains, compared to 61% of the repeats in WT (Figure 4E). Analysis of individual DMTs showed that recovery of IDA *b* in the SNAP-tagged strains could be observed on DMT2 to DMT8, with the C-SNAP closest to WT and M-SNAP showing more DMT-specific variability, reaching almost WT levels on DMTs 6 and 7, but significantly less on the remaining DMTs (Figure 4F).

To determine how the MBO2 polypeptide might be arranged relative to the structures associated with IDA *b*, we treated axonemes with biotin-streptavidin gold (∼1.4 nm) to label the SNAP tags for *in situ* visualization by cryo-ET. Control experiments using silver enhancement procedures to stain samples on gels (Song et al., 2015) confirmed that the SNAP-tags were accessible to streptavidin-gold (Figure 4G). To identify additional densities, i.e., density not visible in WT averages and thus corresponding to the streptavidin-gold labels, we classified the subtomographic volumes of the 96 nm repeats, revealing the tag-density at distinct locations within the repeat (Figure 4H, Figure 5A-F). Specifically, in the N-SNAP rescued strain, the recovery of IDA *b* was evident in 41% of the averaged repeats compared to 61% in WT (Figure 4E), and an additional label-density was observed close to the IDA *b* tail domain in ∼20% of the repeats (compare white and yellow arrowheads in Figure 5A, B). In the M-SNAP strain, 49% of the repeats contained IDA *b* (Figure 4E), and an additional label-density was observed near the distal end of the MBO2-associated complex in ∼15% of the repeats (yellow in Figure 5D). In the C-SNAP strain, 35% of the repeats contained IDA *b* (Figure 4E), and an additional label-density was observed close to the IDA *b* tail and inner junction inner in ∼31% of the repeats) (yellow in Figure 5F). These observations suggest that both the N-terminus and C-terminus of MBO2 are located between RS1 and RS2, in the vicinity of the IDA *b* attachment site, possibly stretching a full 96nm repeat (see Figure 7 and Discussion).

**Figure 5.**
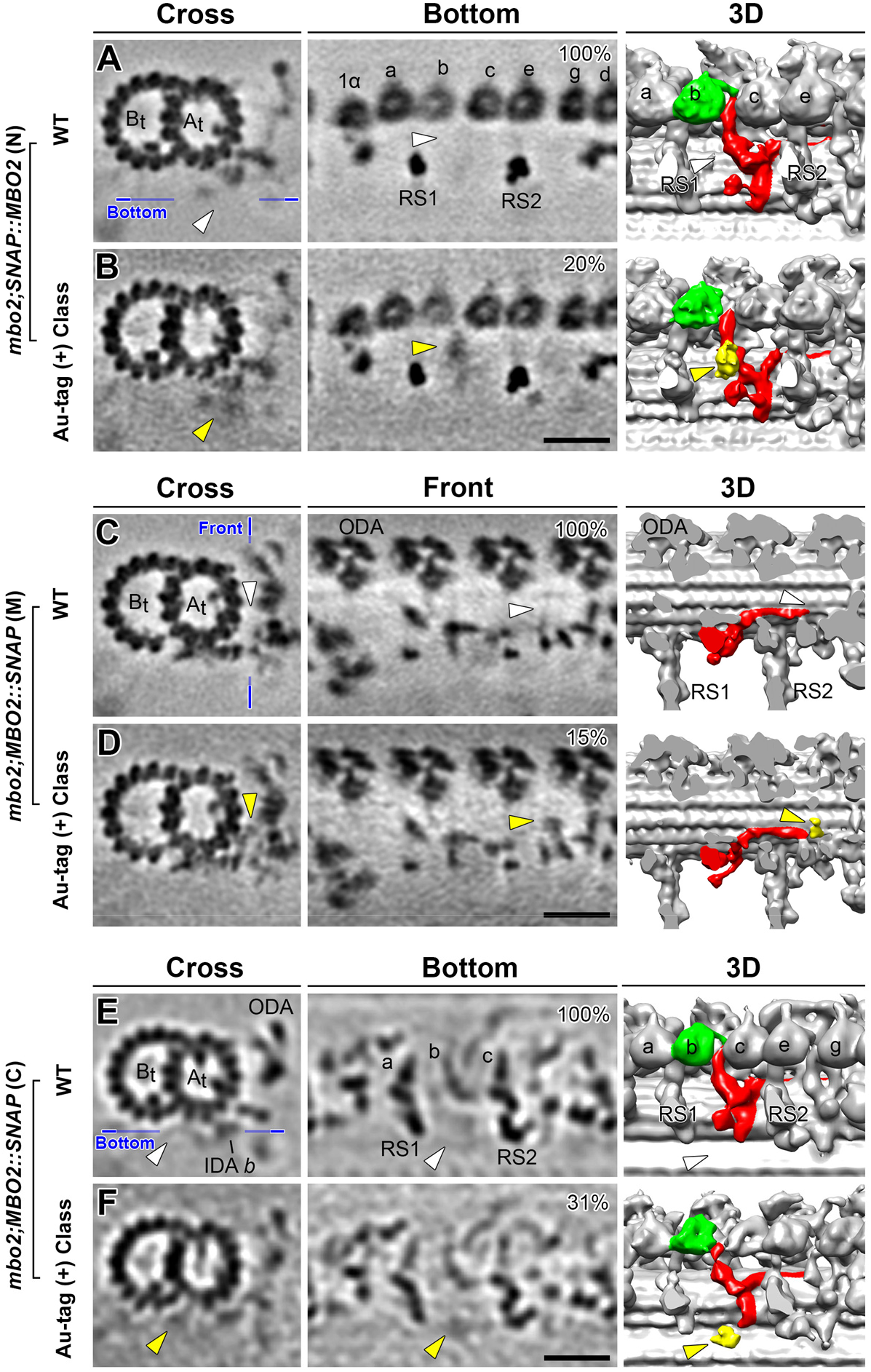
Localization of the N-terminus, middle region, and C-terminus of the MBO2 polypeptide in the 96nm axoneme repeat. The locations of the three different SNAP tags were revealed by comparing WT (**A, C, E**) with streptavidin-gold labeling of different SNAP-tagged rescued strains (**B, D, F**) and class averaging of the axonemal repeats with tag-density. Shown are tomographic slices (left two columns) and 3D isosurface renderings (third column). (**A, B**) The region containing the structures associated with the tail domain of IDA *b* are viewed in cross-sectional and bottom orientations in WT (**A**) and mbo2; *N-SNAP::MBO2* rescued axonemes (**B**). The blue line in (A) indicates the location of the slices viewed in the bottom orientations. The IDA *b* motor domain is colored in green, and other structures that are missing in *mbo2* but were restored in the *N-SNAP::MBO2* rescued axonemes are colored in red. Classification analysis revealed an additional density near the tail domain of IDA *b* in ∼20% of the repeats (indicated by yellow arrowheads in B). WT repeats lack similar density in this position (white arrowheads in A). (**C, D**) Tomographic slices viewed in cross-sectional and front orientations and cropped 3D isosurface renderings viewed in front orientation of the averaged repeats from WT (**C**) and *mbo2; MBO2::M-SNAP* rescued axonemes (**D**). The blue line in (C) indicates the location of the slice viewed in front orientation. **(D)** Classification analysis of *mbo2; MBO2::SNAP* (M) averages revealed an additional density (yellow arrowheads) in ∼15% of the repeats; this density is located on the surface of the A-tubule between protofilaments A04 and A05. A similar density in this was not observed by classification of the WT repeats (white arrowhead in C). (**E, F**) Tomographic slices and 3D isosurface renderings viewed in cross and bottom orientations from WT and *mbo2; MBO2::C-SNAP* axonemes in the region close to the surface of the DMT below IDA *b*. The blue line in (E) indicates the locations of the slices viewed in the bottom orientations in (E) and (F). (**F**) Classification analysis of the *mbo2; MBO::C-SNAP* rescued axonemes revealed an additional density (yellow arrowheads) in ∼31% of the repeats. This density indicates the likely location of the C-terminus of MBO2 close to the inner junction. A similar density was not seen by classification of WT axonemes (white arrowheads in E). Other labels: At/Bt, A- and B-tubule; a-c, single-headed IDAs; ODA, outer dynein arm; RS, radial spoke. Scale bars in A, C, E, 20 nm (valid for all EM images).

**Figure 7.**
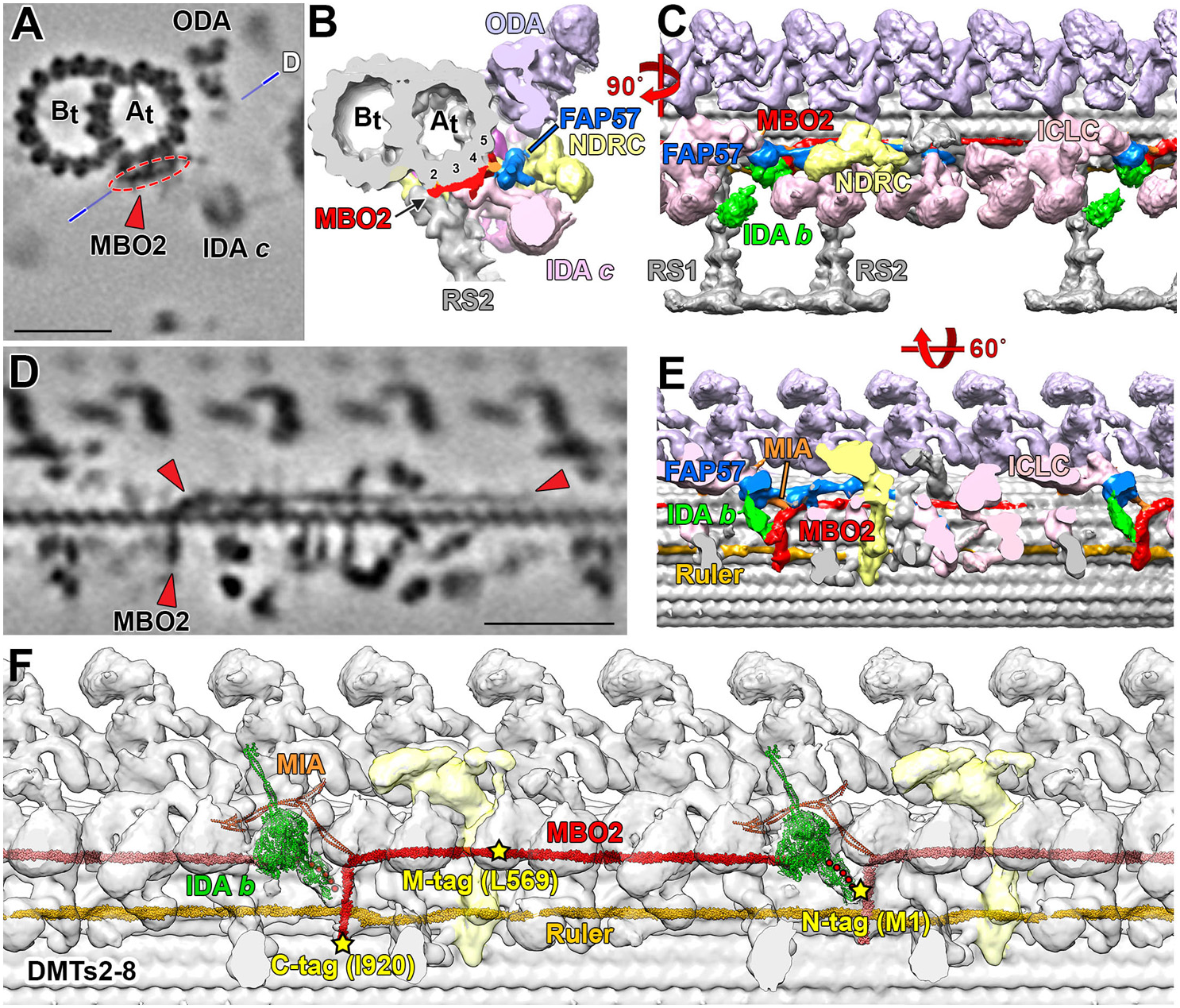
Summary images illustrating the proposed location of the MBO2 complex and its interactions with other regulatory complexes in the 96nm repeat. (**A, B**) Tomographic slice and 3D isosurface rendering of the DMT in cross-section in the region between RS1 and RS2 indicating a filamentous structure on the surface of DMT (circled in red in A and colored in red in B) running from the A02 protofilament to the cleft between protofilaments A04 and A05. (**C**) Rotating the isosurface model 90° along y-axis shows the position of the red structure below IDA *b* (in green) and the globular domain of FAP57 (in blue). The red structure continues along the surface of the DMT, running behind the N-DRC (yellow) and extending into the next 96nm repeat. (**D**) A tomographic slice taken through the DMT along the plane illustrated by the line in (A) and (**E**) the corresponding isosurface model tilted 60 degrees along x-axis from the view in (C) shows the MBO2-associated, L-shaped structure running vertically from protofilament A02 to A04 and then bending and extending distally along the cleft between protofilaments A04 and A05 (indicated by red arrowheads in D and red structure in E). Clipping the isosurface rendering reveals the interactions of the MBO2 complex with the axoneme ruler (gold filamentous structure) located between protofilaments A02 and A03. The MBO2-associated, L-shaped structure extends upward near the base of IDA *b* (green) and the MIA complex (orange) and then turns distally, running below the FAP57 complex and N-DRC and beyond RS2 and RS3S (unshaded) into the next 96 nm repeat, where it fades from view just below the I1 dynein. (**F**) A model for the proposed location of a MBO2/FAP58 heterodimer within the 96 nm repeat. We used AlphaFold2 to predict the structure of a MBO2/FAP58 heterodimer based on the Clustal W alignment of the MBO2, FAP58, and FAP189 polypeptide sequences (Figure S6). We then placed the MBO2 structure in the 96nm repeat based on locations of the SNAP tags detected by streptavidin gold labeling (Figure 5). Because the N-terminal region of MBO2 is disordered, we could not follow the filamentous structure shown in D and E to the site predicted for N-terminal SNAP tag of MBO2, and so we have indicated this flexible region (∼5.5 nm) by a series of red dots. The proposed location for the MBO2/FAP58 heterodimer also agrees with the positioning of a FAP189 homodimer based on mapping residues 689-719 near the MIA complex and residues 384-480 near the FAP78 distal protrusion (Walton et al., 2023). We also added the AlphaFold2 models for the MIA complex and IDA *b* to illustrate how they are proposed to interact with the MBO2/FAP58 heterodimer (Yamamoto et a., 2013; Walton et al., 2023). Scale bars in (A) and (D) are 25 nm.

We did not detect additional densities for streptavidin gold in all repeats of the SNAP-rescues, even though we observed significant recovery of IDA *b* and MBO2-associated structures in all three SNAP-tagged strains (Figure 4E, F). Given the differences between DMTs in the *mbo2* mutant, we considered two possibilities. The first is that the accessibility of the SNAP-tags in the MBO2 subunit to the streptavidin-gold particles might vary within the complex of associated proteins in the axoneme. The second is that our ability to detect the SNAP tags might vary between different DMTs. We therefore checked the tomograms to determine which DMTs were associated with the label densities (Figure 4H). We found that the C-terminal tag was detected on mostly on DMT2 to DMT8. However, the N-terminal and M-SNAP tags were detected more frequently on DMT6 to DMT8 than other DMTs. These observations suggest that the MBO2 subunit is located on DMT2 to DMT8, but the N- and M-SNAP tags are more accessible to gold labeling on DMT6 to DMT8.

## Discussion

### Characteristics of proteins missing or reduced in mbo axonemes and their potential interactions

MBO2 is a conserved, coiled coil protein that is tightly bound along the length of the axoneme, and 2D-PAGE previously indicated that *mbo* axonemes lack eight other proteins (∼33-245 kD) (Segal et al., 1984; Tam and Lefebvre, 2002). Here we used five rounds of isobaric labeling (iTRAQ or TMT) and quantitative MS/MS to identify at least 19 proteins that were missing or reduced in *mbo1* and *mbo2* axonemes and restored in *MBO2-HA* rescued axonemes (Table 1, Supplemental Tables S4, S5). We consider the ten polypeptides that were significantly reduced in the four iTRAQ experiments (using four biological replicates) and the single TMT experiment (with three biological replicates) as the best candidates to be part of the core MBO2 complex (Table 1). The remaining nine proteins that varied in statistical significance between the two different methods may include *bona fide* MBO2 subunits, MBO2-associated proteins present in sub-stoichiometric amounts, subunits of other complexes that interact with MBO2, and other proteins of unknown significance. To gain further insight into these proteins, we analyzed their susceptibility to extraction with high salt (0.6M NaCl/KCl), their co-fractionation with IDAs by FPLC, and their relative abundance in other motility mutants that alter waveform asymmetry or ciliary length. We divided the 19 proteins into five groups as summarized in Table 2 and described below. Several other low abundance proteins were identified as reduced in *mbo2* and restored in *MBO2-HA* in the TMT proteome, and their characteristics are shown in Supplemental Table S4)

The first group contains MBO2 and two other proteins, FAP58 and FAP51, that may dimerize with MBO2 to form the core of the L-shaped structure in the 96 nm axoneme repeat. All three proteins are relatively abundant, coiled coil proteins, resistant to high salt extraction, significantly reduced below ∼25% in *mbo* proteomics experiments, but not significantly altered in *pf12/pacrg, ida8/fap57*, and short mutant axonemes (Figure 1, Tables 1, 2, Supplemental Table S4, Pazour et al., 2005; Lin et al., 2019; Sakato-Antoku and King, 2022). FAP51 is not highly conserved, but MBO2 and FAP58 share significant sequence homology (33.5% identity, Blast P score 3e-44) and even greater homology to different proteins in other species with motile cilia and flagella (Nevers et al., 2017). FAP58 is also closely related (∼80% identity) to a *Chlamydomonas* paralog, FAP189. FAP58, however, is reduced in all mutants with a *mbo* phenotype, whereas FAP189 is unchanged or possibly elevated in *mbo* strains (Table 1, Supplemental Table S4). Thus, the FAP58 and FAP189 paralogs may play distinct but complementary functions in *Chlamydomonas*. A recent cryo-EM single particle analysis visualized parts of the L-shaped structure on the A-tubule with higher resolution, revealing a coiled coil filament. The authors proposed a pseudo-atomic model, placing a FAP189 homodimer within the L-shaped coiled coil structure on the surface of the DMT (Walton et al., 2023). We suggest that MBO2 and FAP58 may form a coiled coil heterodimer that complements the role of a FAP189 homodimer on different DMTs or regions. MBO2 is the orthologue of the vertebrate polypeptide CCDC146 (29.5% identity, Blast P score 1e-103), and *ccdc146* mutations result in defective sperm formation and male infertility (Muroňová et al., 2023). Interestingly, there is only a single orthologue of FAP58 in most species (Nevers, et al., 2017), known as CFAP58/CCDC147 in vertebrates (∼50% identity). CCDC147 is abundantly expressed in the testis and to a lesser extent in the lung, heart, and endometrium, and *ccdc147* mutations are associated with abnormal sperm flagella morphology and rare germline mutations associated with lung cancer (He et al., 2020; Sha et al., 2021; Liu et al., 2016). How CCDC146 and CCDC147 might interact in other species and how *ccdc146* or *ccdc147* mutations affect ciliary waveforms are both unknown.

To identify polypeptides that might facilitate interactions between DHC5 and the MBO2 complex, we searched potential subunits for proteins that were enriched in high salt extracts, reduced in *pf12/pacrg* and/or short mutant axonemes, but not altered in *ida8/fap57* axonemes (Figure 1, Supplemental Table S4, Pazour et al., 2005). This second group of MBO2-associated proteins contains DHC5 and three other proteins with interesting characteristics. FAP343 (∼70 kD) is a coiled coil, WW repeat domain protein with a small region of limited homology to CEP104 (Table 1); it is reduced ∼50% in most *mbo* proteomics experiments. FAP324 (∼70 KD) is significantly reduced below 17% in all *mbo* proteomics experiments and contains an N-terminal coiled coil domain and C-terminal Kelch and FN3 domains. It shares limited homology to KLHDC3, which is abundantly expressed in the testis (Table 2). FAP238 is a small (42 kD), coiled coil protein with an EF-hand domain that may function as a calcium sensor. These three subunits should be further studied to see if they link the base of DHC5/IDA *b* to the L-shaped structure.

The third group of potential MBO2-associated polypeptides contains two proteins, FAP331 and Cre07.g313850, that are conserved paralogs of FAP57 and FAP337 respectively. They are enriched in high salt extracts and reduced in *mbo* and *pf12/pacrg* axonemes (Supplemental Table S4) but increased in *ida8/bop2/fap57* axonemes (Lin et al., 2019; Bustamante-Marin et al., 2020). We re-analyzed the MBO2-associated proteins in *ida8/fap57* axonemes and the FAP57-associated proteins in *mbo* and *pf12/parcg* axonemes (Supplemental Table S4). FAP57 was unchanged, but the *mbo* and *pf12/parcg* mutations appeared to de-stabilize the FAP57 paralogs (FAP331 and FBB7); these may vary on different DMTs or between the proximal and distal regions of the axoneme (Lin et al., 2019).

Comparative proteomics identified a fourth group containing four other subunits (Cre07.g354650, Cre16.g 682850, FAP258, and FAP271) that were reduced in both *mbo* and *pf12/pacrg* (Table 2, Supplemental Table S4). These are less well conserved, mostly coiled proteins that vary widely in size and relative abundance, and there is limited information about their biochemical properties. One subunit, FAP271, shares a sperm-tail PG-repeat domain with STPG1, another protein is abundant in testis. Both Cre16.g682850 (∼87 kD) and FAP271 (∼40 kD) were significantly reduced in all *mbo* proteomes and may localize with MBO2 in the L-shaped structure (Supplemental Table S4).

The last group contains several large (∼101-348 kD), mostly coiled coil proteins that were less significantly reduced in *mbo2*, not altered in *pf12/pacrg* or *ida8/fap57*, but increased in short axonemes (Tables 1, 2, Supplemental Table S4). Most are also poorly conserved, and some are sub-stoichiometric to MBO2 in WT axonemes (Table 2, Pazour et al., 2005; Sakato-Antoku and King, 2023). One exception is FAP308, which is significantly reduced in all *mbo* proteomes and enriched nearly 3-fold in short axonemes, suggesting a potential function in the proximal region of the axoneme (Hwang et al., submitted).

### Cryo-ET of mbo2 reveals DMT specific defects in IDA b and other structures

Cryo-ET of *mbo2* axonemes confirmed the loss of IDA *b* in the 96 nm repeat (Figure 2), and transformation of *mbo2* with tagged *MBO2* constructs rescued the *mbo* motility phenotype and restored assembly of DHC5 and IDA *b* into the axoneme (Figures 1, 4). Thus, although MBO2 and DHC5 are members of distinct biochemical subcomplexes, MBO2-associated proteins are required for stabilizing the assembly of DHC5 and for targeting IDA *b* to a specific site in the 96 nm repeat. To investigate how these complexes might interact within the axoneme, we turned to DMT-specific averaging. Immunofluorescence microscopy first demonstrated that IDA *b* is severely reduced in the proximal 2 μm of WT axonemes (Yagi et al., 2009), and cryo-ET later showed that IDA *b* is also missing or reduced on DMT1, DMT9 and DMT5 in the medial-distal region of WT axonemes (Bui et al., 2012; Lin et al., 2012; 2019; Dymek et al., 2019). In *mbo2*, IDA *b* is reduced on all DMTs (Figures 2, 3) but reassembled on DMT2 to DMT8 in *MBO2* rescued strains (Figure 4). DMT-specific averaging further revealed that the L-shaped structures located below the tail domain of IDA *b* are heterogenous in appearance on different DMTs in both WT and *mbo2* (Figure 3). More specifically, the L-shaped structure (indicated in red in Figures 3,5,6), located between RS1 and RS2 on DMTs 5 to 8 in WT axonemes is missing on DMTs 5 to 8 in *mbo2*. A slightly different L-shaped structure is present on DMTs 2 to 4 in both strains (indicated in purple in Figures 3,5,6). The heterogeneity in structures suggests that some of the proteins reduced in *mbo2* are located on DMTs 2 to 8, whereas other subunits may be restricted to specific regions of the axoneme or a smaller subset of DMTs.

The *mbo*, *pacrg*, and *fap20* mutants also share defects in the assembly of the B-tubule beak-MIP structures located inside DMTs in the proximal region of the axoneme (Supplemental Figure S4; Segal et al., 1984, Dymek et al., 2019; Yanagisawa et al, 2014). More specifically, *mbo2* lacks beaks in DMT5 (Supplemental Figure S4), and both *pacrg* and *fap20* display beak defects in DMT1, 5, and 6 (Yanagisawa et al., 2014; Dymek et al. 2019). The FAP20 and PACRG subunits form the inner junction (IJ) between A- and B-tubules of the DMT (Yanagisawa et al., 2014; Dymek et al., 2019). Some of the overlapping proteins reduced in *mbo2* and *pf12/pacrg* may be involved directly or indirectly in assembly of the beak structures. How defects at the IJ of the DMT (in the case of *fap20* and *parcg*) or on the surface of the A-tubule (in the case of *mbo2*) are both associated with defects in beak structures in the proximal B-tubules *and* DMT-specific defects in assembly of IDA *b* along the length of the axoneme is not yet understood.

Cryo-EM single particle analysis of WT axonemes has recently provided some new insights into the identity of the proteins that form the DMT specific beak structures in *Chlamydomonas* (Leung et al., 2023). These include a group of SAXO proteins that bind to protofilaments B04-B06 and are associated with short tektin filaments in the proximal region of the axoneme. Because the specific SAXO proteins in the B-tubules were not identified (Leung et al. 2023), we re-evaluated the *mbo* and *pf12* proteomes for potential changes in tektin levels. Tektin was reduced ∼25% in *pf12/pacrg* axonemes but was more variable in *mbo* axonemes (Table 1, Supplemental Table S4). Identification and characterization of the B-tubule SAXO proteins and a tektin mutant will be needed to better understand the factors that contribute to the assembly of the beak-MIP structures. The *fap20* and *pf12/pacrg* phenotypes suggest that holes at the IJ of the DMT may influence the stability of tektin and possibly other proteins inside the B-tubule.

Further characterization of missing proteins that are shared between *parcg/pf12, fap20*, and *mbo* mutants could provide new insights into similarities between the strains (i.e., symmetric waveforms, defective beaks, reduced IDA *b*) and overlapping protein network components. Likewise, further characterization of proteins that are uniquely missing in *mbo* mutants could provide a better understanding of the mechanism that converts the ciliary waveform from forwards to backwards swimming.

### Epitope tagged MBO2 constructs rescue the mbo phenotype and most of the DHC5/IDA b defects

Transformation with tagged *MBO2* constructs restores forward swimming, but none of the rescued strains swim at completely WT velocities (Figures 1, 4 and Tam and Lefebvre, 2002). To better understand why swimming velocities might be reduced, we analyzed the re-assemby of DHC5/IDA *b* using multiple proteomic approaches and cryo-ET. In general, the recovery of the swimming velocity could be correlated with the extent of DHC5/IDA *b* recovery (Figure 4, Supplemental Figure S2, Supplemental Table S4). However, disruption of the DHC5 motor domain by itself had negligible effects on motility (Supplemental Figure S3). These observations suggest that the interaction of DHC5 tail domain with the MBO2-associated subunits is more critical for regulating motility than the activity of one IDA motor domain alone. Interactions within the MBO2 complex may influence coordination with other dyneins in the axoneme and/or alter mechanical properties such as resistance or elasticity. Indeed, interactions between dynein motor and tail domains has been shown to alter motor activity in many systems (Rao et al., 2021; McKenney, 2019).

### Localization of the MBO2 SNAP tags within the 96 nm repeat

Given the heterogeneity of structural defects in the *mbo2* mutant, we turned to streptavidin gold labeling of SNAP tags, DMT specific averaging, and class averaging to determine the approximate location of the MBO2 polypeptide. Labeling of the C-terminal SNAP tag and DMT specific averaging suggested that MBO2 is present on DMT2 – DMT8 (Figure 4H). Classification averaging of all DMTs from C-SNAP tagged axonemes detected an additional density close to the inner junction, below the shorter part of the L-shaped structure that is missing from DMT5 to DMT8 in *mbo2* (Figures 5, 6). The additional density marking the N-terminal SNAP tag is located above and slightly proximal of the C-terminal tag. It sits above the surface of the DMT, close to the tail domain of IDA *b*, but below the motor domain (Figures. 5, 6). The additional density marking the M-SNAP tag at amino acid 569 was detected beyond RS2, and near the surface of the DMT, between protofilaments A04 and A05 (Figures 5, 6). This site is close to the distal end of the red structures detected as missing by cryo-ET of the *mbo2* mutant.

The MBO2 sequence is predicted to form several coiled coil domains separated by regions whose structure cannot be predicted with high confidence (Figures 1A, 4A and Tam and Lefebvre, 2002). One possible interpretation based on both the locations of the labels and the size and predicted structure of MBO2 polypeptide is that the MBO2 subunit adopts an extended confirmation spanning the 96 nm repeat, like that observed for the CCDC39/CCDC40 (FAP59/ FAP172) subunits that determine the positions of the radial spokes (Oda et al., 2014; Gui et al., 2021). An alternative interpretation is that the N-terminus of MBO2 is located close to IDA *b* and the polypeptide extends distally towards RS2; the MBO2 subunit then folds back proximally and turns downward towards the C-terminus (Figure 6), so that the N- and C-termini of one MBO2 subunit are both located near the same IDA *b* docking site. We have tried to distinguish between these two possible scenarios as described below.

**Figure 6.**
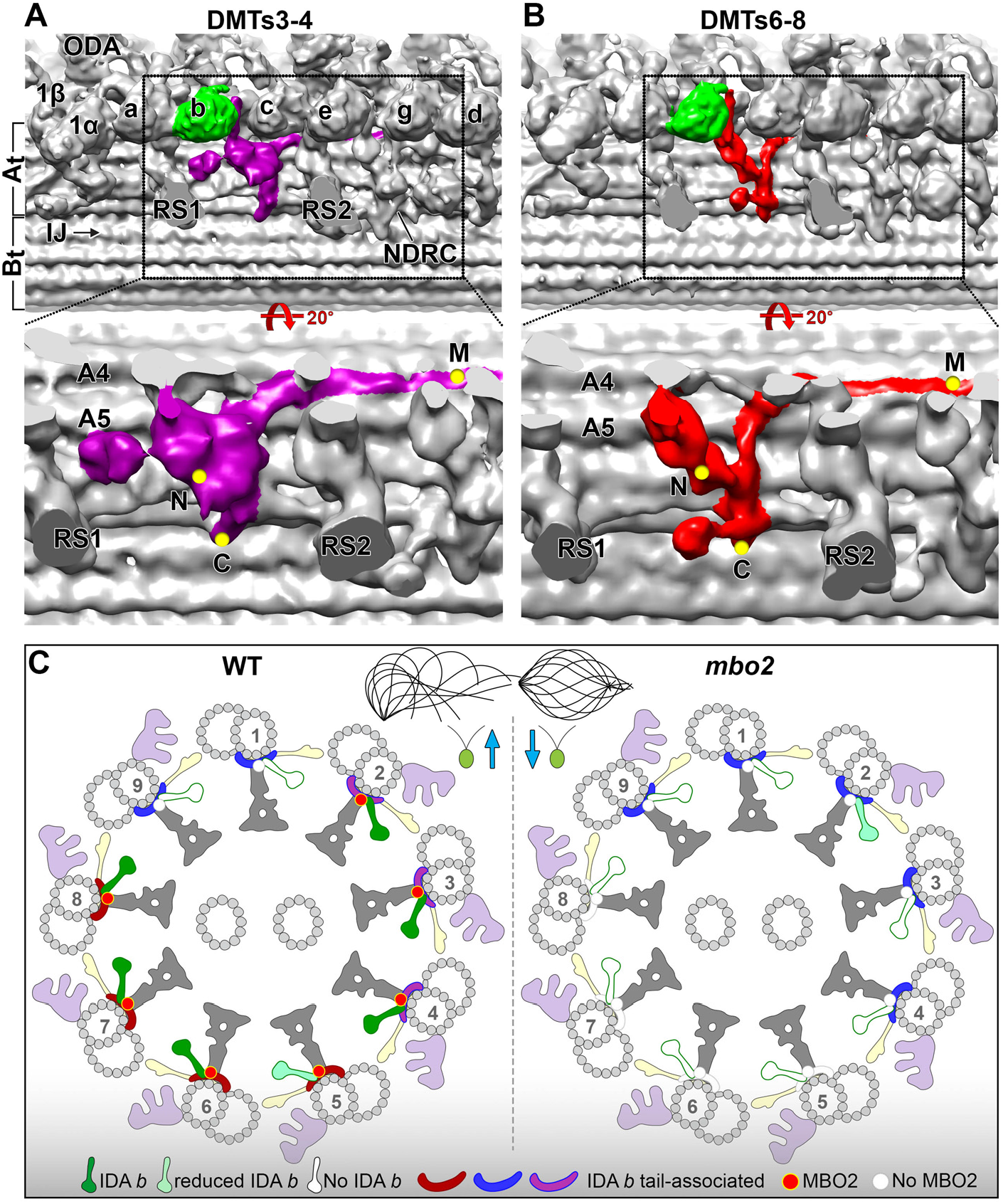
Doublet specific asymmetry of IDA *b* tail-associated structures. **(A, B)** 3D isosurface renderings of the 96-nm repeats obtained by averaging DMTs3-4 (A) and DMTs6-8 (B) of WT axonemes. The regions marked by dashed boxes on the images in the top row were rotated and enlarged in the images shown in the bottom row for better visualization of the structures associated with the IDA *b* tail domain. These structures are highlighted by purple and red colors in DMTs3-4 and DMTs6-8, respectively. Locations of the additional densities seen by gold labeling of the C-terminal, N-terminal, and Mid-region SNAP tags in MBO2 are indicated by the yellow dots. **(C)** Schematic drawings of the cross-section of WT (left) and *mbo2* (right) axonemes showing the arrangement of MBO2 and IDA *b* tail-associated structures across the nine DMTs. Our data suggest that in WT, MBO2 is missing on both DMT1 and 9, because no defects were observed on these DMTs in the *mbo2* mutant. The distinct variations observed in the IDA *b* tail-associated structures on different DMTs are indicated by different colors (green, light green, white). The structure is unchanged between WT and *mbo2* on DMT1 or DMT9 (see blue density) and only slightly affected on DMT2 (purple to fushia). Loss of MBO2 causes the IDA *b* tail-associated structures to be completely missing on DMT5 - DMT8 (white densities) and alters their morphology on DMT3 - DMT4 (from purple to blue), resembling the structures seen on DMT1 and DMT9. Other labels: At/Bt, A- and B-tubule; a-g, single-headed IDAs; 1α, 1β, the I1 dynein α- and β-head; IJ, inner junction; N-DRC, nexin-dynein regulatory complex; ODA, outer dynein arm; RS1, RS2, radial spoke 1 and 2.

### Model for the proposed location of the MBO2 subunit and its interaction with other complexes in the axoneme

We have identified a candidate L-shaped structure in our highest resolution images of WT *Chlamydomonas* axonemes that we propose as the likely location of the MBO2 polypeptide on DMTs 2-8 (Figure 7, Supplemental Table S5). We suggest that the C-terminal region of MBO2 corresponds to a filamentous structure that can be seen close to the surface of the A-tubule in cross-section, with the C-terminus located near protofilament A02 (Figure 7A, B), consistent with the location of the C-terminal SNAP tag shown in Figure 5. The C-terminal region of the filament wraps around the A-tubule (circled in red/colored red in Figure 7A, B) to a cleft between protofilaments A04 and A05. Rotating the images 90° along the y-axis (Figure 7C, D) shows the position of filament running vertically from protofilament A02 to A04 (red arrowheads in Figure 7D) and then turning distally and running along the surface of the DMT between protofilaments A04 and A05 to the red arrowhead on the right (Figure 7D). This filament is modeled in red in the isosurface rendering (Figure 7C). Tilting the isosurface rendering 60° as shown in Figure 7E, the MBO2-associated, L-shaped structure (red) interacts with the axoneme ruler (gold) at its C-terminus and then extends vertically, running below the base of IDA *b* (green) to contact the MIA complex (orange). The red structure turns distally to extend along the surface of the DMT, running below the FAP57 complex (blue) and N-DRC (yellow) and beyond the RS3S (unshaded) into the next 96nm repeat, where it fades just below the I1 dynein, ∼5.5 nm from the predicted location of the N-terminal SNAP tag.

The L-shaped structures present in WT and missing in *mbo2* (Figures 3, 5-7) that we propose as the location of MBO2 are similar in location and organization to a L-shaped, coiled coil structure described in cryo-EM studies of WT axonemes (see Figure 1 in Ma et al., 2019; Extended Data Figure 4 in Gui et al., 2021). This coiled coil structure has recently been proposed as the location of FAP189 based on limited fitting of short regions of an AlphaFold2 predicted structure for a FAP189 homodimer (predicted in several sequence fragments) to a single particle cryo-EM reconstruction of WT axonemes (Extended Data Figures 5 and 8 in Walton et al., 2023). However, we have found that the decrease in MBO2 seen in all the *mbo* mutants is more closely correlated with a decrease in FAP58, whereas FAP189 levels are essentially unchanged or even increased in the *mbo* mutant axonemes (Table 1, Supplemental Tables S4). This suggests that at least on DMTs 5-8, where the L-shaped structure is missing in *mbo2*, MBO2 and not FAP189 form the L-shaped coiled coil filament. We predict that MBO2 and FAP58 form a heterodimer that serves as a scaffold for the assembly of other polypeptides missing in *mbo* axonemes. We further hypothesize that the MBO2/FAP58 heterodimer and FAP189 homodimer perform distinct but complementary functions in stabilizing the binding of other proteins in the 96 nm repeat. MBO2/FAP58 and FAP189 may occupy similar sites on different DMTs or vary in proximal, medial, or distal regions. For instance, a FAP189 homodimer may form part of the L-shaped densities that remain on other DMTs in the *mbo2* mutant. The FAP58 and FAP189 subunits may also partially compensate for one another in different mutants such as *mbo1* where FAP189 appears to be elevated, similar to that proposed for different paralogs of the FAP57/FAP337 complex in *fap57* mutants (Lin et al., 2019). Studies of a *fap189* mutant in *Chlamydomonas* will be required to resolve these questions.

To test the hypothesis that MBO2 and FAP58 form a heterodimer, we aligned the polypeptide sequences for MBO2, FAP58, and FAP189 using Clustal W (Supplementary Fig. S6). We then used AlphaFold2 (Mirata et al.2022) to predict a potential MBO2/FAP58 heterodimer structure using full length sequences. The resulting structure prediction shows extensive coiled coil domains separated by short, unstructured regions and slightly longer, unstructured regions at the N- and C-termini of both subunits. We placed the Alphafold2 predicted structure of the MBO2/FAP58 heterodimer within the 96 nm repeat based on the positions of the MBO2-SNAP tags detected by streptavidin gold labeling (Figures 5, 6). Because the N-terminal region of MBO2 is thought to be disordered, we could not follow the filamentous structure completely to a site near the IDA *b* tail domain that is predicted to be the N-terminus of MBO2 (Figures 5, 7D), and so this region is represented by a series of red dots (Figure 7E). However, the positions of the M-SNAP tag at leucine 569 and the C-SNAP tag at isoleucine 920 are consistent with the lengths of the predicted coiled coil domains (Figures 5, 7E). The placement of the MBO2/FAP58 heterodimer is also consistent with the mapped positions of short segments of amino acids that are conserved between FAP189 and FAP58 (See Figure Legend 7 and Walton et al., 2023).

FAP189 has been associated with subunits of the MIA complex and FAP57 complex based on chemical cross-linking and immunoprecipitation experiments (Yamamoto et al., 2013). Given the extensive sequence homology between FAP189 and FAP58 (∼80% sequence identity), it is likely the MBO2/FAP58 heterodimer also interacts with both the MIA complex and the FAP57 complex (Figure 7E, Supplemental Figure S7), which is consistent with the proximity of these structures observed by cryo-ET. In *Chlamydomonas*, *fap57* mutants are associated with reductions in FAP57 and increases in two FAP57 paralogs (FBB7 and FAP331). The *fap57* mutants also display DMT specific defects in assembly of IDA *g* and *d*. We proposed that the increases in the levels of the FAP57 paralogs could partially compensate for the loss of FAP57 on certain DMTs in the *fap57* mutant (Lin et al., 2019). Interestingly, we have not seen any evidence that the *mbo* mutants disrupt the assembly of the MIA complex or FAP57 (Table 1, Supplemental Table S4). However, we have detected reductions in the levels of two minor FAP57 paralogs, FAP331 and FBB7, in the *mbo* strains (Tables 1, 2, Supplemental Table S4). FAP57 and its paralogs also interact with two highly conserved, EF-hand, WD repeat proteins known as FAP337 and Cre07.g313850 (Lin et al., 2019), and both FAP337 and Cre07.g313850 are variably reduced in *mbo* mutants (Tables 1, 2, Supplemental Table S4). Studies in *Tetrahymena* using chemical cross-linking, single particle cryo-EM and AlphaFold2 modeling have recently localized FAP337 to a “staple” formed by the coiled-coil heterodimer of CCDC96/CCDC113 (FAP184/FAP263) that is located distal to the N-DRC (Bazan et al., 2021; Ghanaeian et al., 2023). FAP337 also interacts with the base of IDA *d* and the coiled-coil domains of two FAP57 paralogs, CFAP57A/CFAP57C, and CFAP57A/C interacts in turn with bases of IDA *d* and *g* (Ghanaeian et al., 2023). Collectively these observations illustrate how FAP57 and FAP337 (and their paralogs) can interconnect the MIA complex, the N-DRC, CCDC96/CCDC113, and stabilize the binding of IDA *d* and *g* (Lin et al., 2019; Bazan et al,.2021; Ghanaeian et al., 2023).

In *Tetrahymena*, the CFAP57 paralogues also interact with an unidentified L-shaped coiled-coil structure like the MBO2/FAP58/FAP189 structure described above (Ghanaeian et al., 2023; Walton et al., 2023). Our observations that FAP331 and Cre07.g313850 are reduced in *mbo* mutants are consistent with an interaction between the C-terminal coiled domains of the FAP57 paralogs and the MBO2-associated L-shaped structure, although the precise site of interaction has not been mapped. We have not identified any defects on the assembly of IDA *d* (DHC2) and IDA *g* (DHC3, DHC7) in the *mbo* strains, but impact of any reductions in FAP331 and Cre07.g313850 may be offset by the WT levels of FAP57 and numerous connections between the other structures in this region. We propose that FAP331 and Cre07.g313850 are located in similar positions as FAP57 and FAP337, and that they also serve a partially redundant and possibly DMT-specific function in *Chlamydomonas.* The proposed location of FAP57 and its paralogs relative to other structures in the 96nm repeat is illustrated in Supplemental Figure S7.

### Interactions of the MBO2 complex with other structures in the axoneme and implications for motility

Previous work revealed that generation of ciliary and flagellar waveforms requires an asymmetric distribution of force generated by dynein arm activity along the cilia (Lin and Nicastro, 2018). MBO2 and its associated structures demonstrate that cilia have intrinsic asymmetry built into the structure of the axoneme, as well as their interconnections to multiple regulatory complexes, to facilitate both mechanical and chemical feedback control on dynein arm activity. A biophysical study of the beating patterns in *Chlamydomonas,* including the *mbo2* mutant, showed that the asymmetric ciliary beating waveform of *Chlamydomonas* WT could be mathematically separated into static and dynamic components. The study concluded that *mbo2* lacks the static component and – correctly - predicted IDA defects in the *mbo2* mutant (Sartori et al., 2016). Our findings suggest that MBO2 and its associated IDA *b* structures could contribute to generate the static component of the ciliary waveform in *Chlamydomonas*.

## Materials and Methods

### Culture conditions, genetic analyses, and strain construction

Strains used in this study (Supplemental Table S1) were maintained on Tris-acetate phosphate (TAP) medium, but occasionally resuspended in liquid minimal medium or 10 mM Hepes, pH 7.6, to facilitate flagellar assembly and mating. Transformants were selected by co-transformation with pSI103 (encoding the *aphVIII* gene) (Sizova et al., 2001) or pHyg2 (encoding the *aphVII* gene) (Berthold et al., 2022) and plating on media containing 10 μg/ml paromomycin or hygromycin B.

### Epitope tagging of MBO2 and characterization of candidate dhc5 mutations

Purification of genomic and plasmid DNA, restriction enzyme digests, agarose gels, and PCR reactions, were performed as previously described (Lin et al., 2019). All primers used for SNAP tagging of MBO2 and characterizing insertions into the *DHC5* gene are listed in Supplemental Table S2. The plasmid containing the wild-type *MBO2* gene tagged with a 2HA tag was previously described (Tam and Lefebvre, 2002), and the predicted polypeptide sequence is shown in Supplemental Figure S1. To make a construct encoding a SNAP tag at the N-terminus of MBO2, a 2033 bp gene fragment spanning two *Apa*I sites and encoding a codon optimized SNAP tag and first three exons and introns of *MBO2* was synthesized and cloned into pUC57 (Genewiz, Azenta Life Sciences, South Plainfield, NJ). This fragment was amplified by PCR with primers spanning the *Apa*I sites and complementary to *MBO2*. After gel purification, the SNAP-tagged fragment was assembled into a *Apa*I digested MBO2-HA plasmid using NEBuilder (New England Biolabs, Ipswich, MA). The final *N-SNAP-MBO2-HA* construct encodes an MBO2 polypeptide with a SNAP tag at its N-terminus and a 2-HA tag located between amino acids 885 and 886 of the original MBO2 sequence (Supplemental Figure S1).

To insert a SNAP tag near the middle of MBO2, a 1839 bp region between two *Kpn*I sites was amplified by PCR and subcloned into pGEM-T Easy (Promega, Madison, WI). This subclone was subjected to site-directed mutagenesis using the Q5 site-directed mutagenesis kit (New England Biolabs) to create a new *Hind*III site at the position encoding amino acid 569. A SNAP tag was amplified from a codon-optimized SNAP plasmid (Song et al., 2015) using primers containing *Hind*III sites and complementary to MBO2, gel purified, and assembled into the *Hind*III site of the *Kpn*I subclone using NEBuilder. The tagged *Kpn*I subclone was then amplified with primers containing *Kpn*I sites, gel purified, and assembled into a *Kpn*I-digested MBO2-HA plasmid. The final *MBO2-M-SNAP-HA* construct encodes an MBO2 polypeptide with a SNAP tag located between amino acids 569 and 570 and a 2-HA tag between amino acids 885 and 886 of the original MBO2 sequence (Supplemental Figure S1).

To tag the C-terminus of MBO2, a 1018 gene fragment spanning two *Hpa*I sites and containing the last exon of MBO2 and codon optimized SNAP tag before the stop codon was synthesized and cloned into pUC57 (Genewiz). This fragment was then amplified using primers containing *Hpa*I sites, gel purified, and assembled into the *Hpa*I digested MBO2-HA plasmid. The final MBO2-C-SNAP construct encodes an MBO2 polypeptide with a SNAP tag at its C-terminus but lacking the HA tag (Supplemental Figure S1).

All new constructs were verified by sequencing (GeneWiz), analyzed using the Sequencher (Gene Codes, Ann Arbor, MI) and MacVector (Apex, NC) software packages, and linearized with *Eco*R1 or *Bam*H1 prior to co-transformation into *mbo2-4*.

### Phase contrast microscopy and measurements of swimming velocity

Motility phenotypes were assessed by phase contrast microscopy using a 20x or 40x objective on a Zeiss Axioskop microscope. Measurements of swimming velocities were made from recordings using a Rolera-MGi EM-CCD camera (Q-imaging, Surrey, BC, Canada) and the Metamorph software (Molecular Devices, San Jose, CA) (VanderWaal et al., 2011; Bower et al., 2013, 2018; Reck et al., 2016). At least 3 independent experiments were performed for each strain. Data are presented as the mean +/− standard deviation using the student’s *t*-test. Images of forward movement were obtained by collecting 1 sec exposures of cells imaged with the 20x objective. Selected images were cropped, rotated, and labeled in Image J and Adobe Photoshop (San Jose, CA).

### Fractionation of axonemes, SDS-PAGE, Western blot, and tandem mass spectrometry (MS/MS) analysis of gel bands

*Chlamydomonas* whole cell lysates, flagella, and axonemes were prepared as previously described (Witman, 1986; Bower et al., 2013, 2018; Reck et al., 2016) using 0.1-1.0% Nonidet-P-40 to remove membrane plus matrix proteins. Purified axonemes were resuspended in HMEEN (10 mM Hepes, pH 7.4, 5 mM MgSO4, 1 mM EGTA, 0.1 mM EDTA, 30 mM NaCl) plus 1 mM DTT and 0.1 μg/ml protease inhibitors (leupeptin, aprotinin, pepstatin), and extracted with HMEEN containing 0.6M NaCl, 0.2M NaI, 0.4M NaI, or 0.6M NaI. Some samples were also extracted with 0.1-0.7% Sarkosyl. The 0.6M extracts from *pf28* (containing IDAs) were diluted 10-fold and fractionated by Mono-Q ion-exchange FPLC chromatography (Gardner et al., 1994). Samples were separated on 5-15% polyacrylamide gradient gels and silver stained or transferred to Immobilon P and probed with different antibodies (Supplemental Figure S2, Supplemental Table S3) (Bower et al., 2013; 2018). A subset of FPLC fractions were analyzed by SDS-PAGE and MS/MS to identify polypeptides that co-fractionate with DHC5. Selected bands were excised and analyzed as previously described (Lin et al., 2019).

Because DHCs vary widely in abundance, purified axonemes were also fractionated by SDS-PAGE, stained briefly with Coomassie blue, and the DHC region was excised from the gel to improve the signal to noise. Following extraction and trypsin digestion, 3-5 replicates per sample were analyzed by MS/MS, and both the total number of peptides and total number of assigned spectra per HC isoform were determined. The relative abundance of each DHC was estimated by spectral counting (Zhu et al., 2010) and expressed as a percentage of the total spectra identified for the 1-alpha and 1-beta DHCs of the I1 dynein as previously described (Bower et al., 2013, 2018). The DHCs were also identified and quantified using the SEQUEST algorithm (Eng et al., 1994) and Proteome Discover 2.3 (Thermo Scientific). For internal calibration of the peptide masses, the re-calibration node was used with a 20 ppm mass tolerance and carbamidomethyl cysteine as a fixed modification. For protein identification, we used the *Chlamydomonas reinhardtii* v5.6 protein FASTA database concatenated with a common lab contaminant database (https://www.thegpm.org/crap/) and the following SEQUEST search parameters: semi-trypsin, 2 missed cleavage sites, minimum peptide length 6, precursor mass tolerance 12 ppm, fragment mass tolerance 0.1 Da, dynamic modifications: oxidation of methionine, deamidation of asparagine and glutamine and pyro-glutamic acid modification of N-terminal glutamine, and carbamidomethyl cysteine as a fixed modification. We used the Percolator algorithm with a concatenated target-decoy database approach to control the false discovery rate (Brosch et al., 2009; Spivak et al., 2009). Each sample was analyzed in triplicate and quantified using the label free quantification workflow that includes steps for feature extraction, chromatographic alignment, peptide mapping to features, protein abundance calculation, normalization, protein relative abundance ratio calculation and hypothesis testing for significance of relative fold change. For each file, we applied the untargeted Minora Feature Detector algorithm, which is similar to the match between runs setting in MaxLFQ (Cox et al., 2014). Peptides were mapped to retention time-aligned consensus features across samples with the requirement that at least one sample contains a peptide spectral match. For each sample, protein abundances were calculated by summing abundances of consensus features for related peptides. We normalized protein abundances for each sample using two proteins: DHC1 (178 distinct peptide sequences) and DHC10 (152 distinct peptide sequences). We set the hypothesis test method to ANOVA and report p-values that were adjusted using the Benjamini-Hochberg correction for false discovery rate (FDR).

### Preparation of samples for iTRAQ or TMT labeling and MS/MS analysis of whole axonemes

iTRAQ labeling: Axonemes were washed in 10 mM Hepes pH 7.4 to remove salt, DTT, and protease inhibitors, then resuspended in 0.5M triethylammonium bicarbonate pH 8.5 and processed for trypsin digestion and iTRAQ labeling as described in detail (Bower et al., 2013, 2018; Reck et al., 2016). Duplicate aliquots of axonemes (50-60 μg each) from each strain were reacted with 4-plex iTRAQ reagents (114-117, AB Sciex, Foster City, CA) to obtain two technical replicates per biological sample. The four labeled aliquots were mixed and processed to remove excess trypsin, unreacted iTRAQ reagents, and buffer. The combined sample (containing two control aliquots with different iTRAQ labels and two mutant aliquots with different iTRAQ labels) was fractionated offline using high pH, C18 reversed phase chromatography (Reck et al., 2016). Approximately 500ng of each peptide fraction was analyzed by LC-MS on a Velos Orbitrap mass spectrometer (Thermo Scientific, Waltham MA). Online capillary LC, MS/MS, database searching, and protein identification were performed as previously described (Lin-Moshier et al., 2013; Reck et al., 2016) using ProteinPilot software version 4.5 or 5.0 (AB Sciex, Foster City, CA) and version 5.5 of the *Chlamydomonas* genome database (https://phytozome.jgi.doe.gov/pz/portal.html). The bias factors for all samples were normalized to alpha and beta tubulin (Reck et al. 2016). The relative amount of protein in each aliquot was compared to that present in the control aliquot to obtain a protein ratio. The WT/WT or HA/HA ratios indicated the variability in labeling and protein loading between technical replicates of the same sample (typically less than 10% for all proteins). Two iTRAQ experiments with independent biological replicates were performed for *mbo1*, *mbo2,* and *pf12.* Between 689 - 919 proteins were identified at a 1% false discovery rate in each experiment. The protein lists were filtered using a minimum of 6 peptides per protein, and those proteins that were significantly reduced (P < 0.05) in all samples using Benjamini - Hochberg correction were further analyzed. We also re-analyzed proteomics data from other iTRAQ experiments using two biological replicates of *ida8* axonemes with defects in the FAP57 complex (Lin et al., 2019), and two biological replicates of *pf9-2; pf28* axonemes, a double mutant that lacks ODAs and I1 dynein and assembles short (∼2.9 μm) flagella (Porter et al., 1992; Hwang et al, submitted).

TMT labeling: The proteins altered in *mbo2* axonemes were analyzed in a third experiment using three independent biological replicates of WT, *mbo2*, and *MBO2-HA* axonemes and TMT16 plex labeling. Samples were digested and labeled with TMT reagents, dried down, and then fractionated offline as described above. Approximately ∼300-600 nanograms of each fraction was analyzed by capillary LC-MS with a Dionex UltiMate 3000 RSLC nano system on-line with an Orbitrap Eclipse mass spectrometer (Thermo Scientific) with FAIMS (high-field asymmetric waveform ion mobility) separation as described by Weise et al., 2023 with minor modifications. The LC profile was 5 to 8% solvent B at 2.5 minutes, 21% B at 135 minutes, 34% B at 180 minutes and 90%B at 182 minutes with a flowrate of 315 nl/minute, where solvent A was 0.1% formic acid in water and solvent B was 0.1% formic acid in ACN. The MS2 settings were: 0.7 Da quadrupole isolation window, 38% fixed collision energy, Orbitrap detection with 50K resolution at 200 *m/z*, first mass fixed at 110 *m/z*, 150 msec max injection time, 250% (1.25 x 10E5) AGC, 45 sec dynamic exclusion duration with +/− 10 ppm mass tolerance and exclusion lists were not shared among compensation voltages.

The peptide output from the tandem MS was processed using SEQUEST algorithm in Proteome Discoverer 2.5. The *Chlamydomonas reinhardtii* protein sequence database was downloaded from https://phytozome-next.jgi.doe.gov/info/Creinhardtii_v5.6 and merged with two additional sequences, DRC1 and FAP58, plus a common lab contaminant protein database (http://www.thegpm.org/cRAP/index.html) (16,631 total protein sequences). The database search parameters were as previously described (Weise et al., 2023) with the exception that the fragment ion tolerance was 0.08 Da. Peptides and proteins were identified using a 1% False Discovery Rate and the Percolator algorithm (Kall et al., 2007) in Proteome Discover and quantified as described in Weise et al, 2023, with the exception that normalization was performed using alpha and beta tubulin. The software used for analysis of TMT labeled samples differs in several ways from the Protein Pilot software used for analysis of iTRAQ labeled samples, but both methods provide estimates of relative protein ratios based on the quantification of isobaric tags.

### Cryo-sample preparation, cryo-electron tomography and image processing

The purified axoneme pellet was resuspended in HMEEK buffer (30 mM HEPES, pH7.4, 5 mM MgSO_4_, 1 mM EGTA, 0.1 mM EDTA, 25 mM KCl), and the suspension was directly used for cryo-sample preparation. For visualization of the SNAP tags, streptavidin nanogold labeling was performed on axonemes from strains rescued with SNAP-tagged versions of the MBO2 as previously described (Song et al., 2015). Briefly, 1 µl of 1 mM BG-(PEG)12-biotin (New England Biolabs; PEG linker available on request) was added to 200 µl of axonemes. A control sample was also prepared without added BG-(PEG)12-biotin. Both suspensions were incubated overnight at 4 °C, followed by three cycles of resuspension with HMEEK buffer and centrifugation at 10,000g for 10 min at 4 °C. The axoneme pellets were resuspended in 200 µl of buffer, and then either 5 µl of 1.4-nm-sized streptavidin nanogold particles (strep-Au, Nanoprobes, Inc) or 5 µl of buffer were added, and the two suspensions were incubated at 4 °C in the dark for 3 h with rotation. The samples were then diluted with 1 ml of HMEEK buffer, pelleted by centrifugation at 10,000 g for 10 min at 4 °C, carefully resuspended in 200 µl of HMEEK buffer, and used for cryo-sample preparation.

Cryo-sample preparation, cryo-ET and image processing were done as previously described (Nicastro et al., 2006; Nicastro, 2009; Heuser et al., 2009; Lin et al., 2014). Briefly, Quantifoil copper grids (Quantifoil Micro Tools, Jena, Germany) with a holey carbon film (R2/2, 200 mesh) were glow discharged for 30 seconds at −40 mA and loaded with 3 µl of axoneme sample and 1 µl of BSA-coated, five-fold concentrated 10 nm colloidal gold (Sigma-Aldrich, St. Louis, MO) (Ivancu et al, 2006). After brief mixing, grids were blotted from the back side with filter paper for 1.5-3 seconds and plunge frozen in liquid ethane using a home-made plunger. Vitrified samples were either loaded onto a cryo holder (Gatan, Inc., Pleasanton, CA) and transferred to a Tecnai F30 or assembled into autogrids and transferred to a Titan Krios transmission electron microscope (ThermoFisher Scientific, Waltham, MA) for imaging. Single-axis tilt series of non-compressed, intact axonemes were acquired using the software package SerialEM (Mastronarde, 2005). Typically, 50 to 70 images were recorded for each tilt series while the specimen was tilted from about −65 to +65° in 1.5 to 2.5° increments. A dose-symmetric scheme was applied for data collection (Hagen et al., 2017). The magnification was set to 13,500X (∼1 nm effective pixel size) with −6 to −8 µm defocus (for Tecnai F30) or 26,000X (∼0.55 nm effective pixel size) with −0.5 µm defocus using a Volta phase plate (Danev et al., 2014) (for Titan Krios). The microscope was operated in low dose mode at 300 keV and the accumulative electron dose of the sample was restricted to ∼100 e/Å^2^ to minimize radiation damage. Electron micrographs were recorded with a 2k x 2k CCD camera (on Tecnai F30) or with a 4k x 4k K2 electron direct camera. The K2 camera was operated in counting mode (0.4 s exposure time per frame and 15 frames per tilt series with a dose rate of 8 e^-^/p/s. Both TEMs were equipped with a post-column energy filter (Gatan, Inc., Pleasanton, CA) that was operated in zero-loss mode with a slit width of 20 eV.

The raw frames from the K2 camera were aligned for motion-correction with a script from the IMOD software (Kremer et al., 1996). 3D tomograms were reconstructed using fiducial alignment of the tilt series images and weighted backprojection using IMOD. Sub-tomograms containing the 96 nm repeat units were further aligned and averaged using PEET (Nicastro et al., 2006), resulting in averaged 3D structures with compensated missing wedge effect, reduced noise and thus increased resolution. For doublet-specific averaging, the nine outer DMTs were identified based on DMT-specific features (Bui *et al*., 2012; Lin *et al*., 2012), and repeats from individual DMTs were averaged. To further analyze structural defects that appeared heterogeneous or to identify the sites labeled with nanogold particles, classification analyses were performed on the aligned sub-tomograms using the PEET program (Heumann *et al*., 2011). Appropriate masks were applied to focus the classification analysis on specific regions of interest, and sub-tomograms containing the same structures were grouped into class averages. The structures were mapped onto their respective locations in the raw tomograms to determine the distribution of the different classes. The numbers of tomograms and sub-tomograms that were analyzed and the resolutions of the resulting averages are summarized in Supplemental Table S5. The resolution was estimated at the center of the sub-tomogram volume using the Fourier shell correlation method with a criterion of 0.5. The structures were visualized as 2D tomographic slices and 3D isosurface renderings using IMOD and UCSF Chimera (Pettersen *et al*., 2004), respectively. The structure of the MBO2/FAP58 heterodimer was predicted using the AlphaFold 2 software (Jumper et al., 2021; Mirdita et al., 2022). To place the pseudo-atomic model into the L-shaped structure, the two longest coiled coil regions were arranged into one long filament and the shorter coiled coil tilted into an about 90° angle relative to the filament (Figure 7F).

### Online supplemental material

Supplemental Figure S1 shows the predicted amino acid sequences of MBO2 and its epitope-tagged variants. Supplemental Figure S2 shows the biochemical characterization of protein samples used for mass spectrometry. Supplemental Figure S3 shows the characterization of the candidate *dhc5* mutants. Supplemental Figure S4 illustrates the defects in the B-tubule beak structures in the *mbo2-4* axonemes. Supplemental Figure S5 shows the Clustal W alignment of the MBO2, FAP58, and FAP189 polypeptide sequences. Supplemental Figure S6 illustrates a model of the proposed interactions of the MBO2 complex with other regulatory complexes in the axoneme.

Supplemental Table S1 lists the strains used in this study. Supplemental Table S2 lists the oligonucleotide primers used for PCR characterization of the candidate *dhc5* mutations and cloning to generate SNAP-tagged *MBO2* constructs. Supplemental Table S3 lists the antibodies used in this study. Supplemental Table S4 lists the protein ratios for the MBO-associated and FAP57-associated proteins obtained identified by iTRAQ labeling and MS/MS analsysis of *mbo1, mbo2, pf12, ida8*, and short *pf9-2; pf28* axonemes, additional protein ratios identified by TMT labeling and MS/MS of *mbo2* and *MBO2-HA* axonemes, polypeptides identified by MS/MS by high salt extraction and FPLC analysis of DHC5. Supplemental Table S5 summarizes the number of tomograms analyzed and the resolution of the averages. References specific to the supplemental material are also included.

## Abbreviations

CSC: Calmodulin- and spoke-associated complex
CP: Central pair
CLiP: *Chlamydomonas* Library Project
DIC: Differential interference contrast
DHC: Dynein heavy chain
ET: Electron tomography
FAP: Flagellar associated polypeptide
HA: Hemagglutinin
IDA: Inner dynein arm
IC: Intermediate chain
IFT: Intraflagellar transport
iTRAQ: Isobaric tag for relative and absolute quantitation
LC: Light chain
MBO: Move backwards only
N-DRC: Nexin-dynein regulatory complex
ODA: Outer dynein arm
*pf*: Paralyzed flagella
PEET: Particle Estimation for Electron Tomography
PCR: Polymerase chain reaction
PCD: Primary ciliary dyskinesia
RS: Radial spoke
RSP: Radial spoke protein
RS3S: Radial spoke 3 stand-in
MS/MS: Tandem mass spectrometry
TMT: Tandem mass tag
TEM: Transmission electron microscopy
TAP: Tris-acetate-phosphate
WT: Wild-type

## Acknowledgements

We thank LeeAnn Higgins, Todd Markowski, Bruce Witthun, and Kevin Murray in the Center for Mass Spectrometry and Proteomics at the University of Minnesota (UMN) for expert assistance with iTRAQ and TMT labeling, mass spectrometry, spectral counting, and data analysis. This center is supported by multiple grants including the National Science Foundation major research instrumentation grants 9871237 and 0215759 as described at https://cbs.umn.edu/cmsp/about. We thank Yanhe Zhao (UT Southwestern) for assistance with the AlphaFold2 predictions of the MBO2/FAP58 heterodimer and FAP189 homodimer. We also thank Matt Laudon and the *Chlamydomonas* Resource Center (https://www.chlamycollection.org/) for strains. This center is supported by the National Science Foundation Living Stock Collections for Biological Research program (NSF grants 0951671 and 00017383). The Porter laboratory also acknowledges the dedicated assistance of multiple UMN undergraduates including Jared Reick, Emma Arnold, and Jad Awada. We also thank Chen Xu (Brandeis University) and Daniel Stoddard (UT Southwestern Medical Center) for dedicated training and maintenance of EM facilities. The UTSW cryo-electron microscope facility is funded in part by a CPRIT Core Facility Award (RP170644). Richard Linck (UMN), Toshiki Yagi (Prefectural University of Hiroshima), Win Sale (Emory University), Ritsu Kamiya (Tokyo University), Paul Lefebvre (UMN), and Pinfen Yang (Marquette University) generously supplied antibodies as listed in Supplemental Table 3. This work was supported by National Institutes of Health grants to M.E.P (GM055667) and D.N. (GM083122). The authors declare no competing financial interests.

## Author contributions

M.E.P and D.N. designed research and obtained funding. G.F. carried out axoneme preparation, cryo-electron tomography, and sub-tomogram averaging. D.T. performed PCR, generated the epitope-tagged constructs, and transformed and screened all mutant strains. J.R., J.S., K.A., and R.B. isolated axonemes, performed SDS-PAGE and Western blots, prepared samples for mass spectrometry, carried out immunofluorescence studies, and measured swimming velocities. M.E.P. analyzed proteomics data and prepared final figures. G.F., L.G., and D.N. analyzed cryo-ET data and prepared final figures. M.E.P., G.F., and D.N. wrote the paper with input from all authors.

## Supplemental Information for The MBO2 complex targets assembly of inner arm dynein *b* and reveals additional doublet microtubule asymmetries

**Figure S1.**
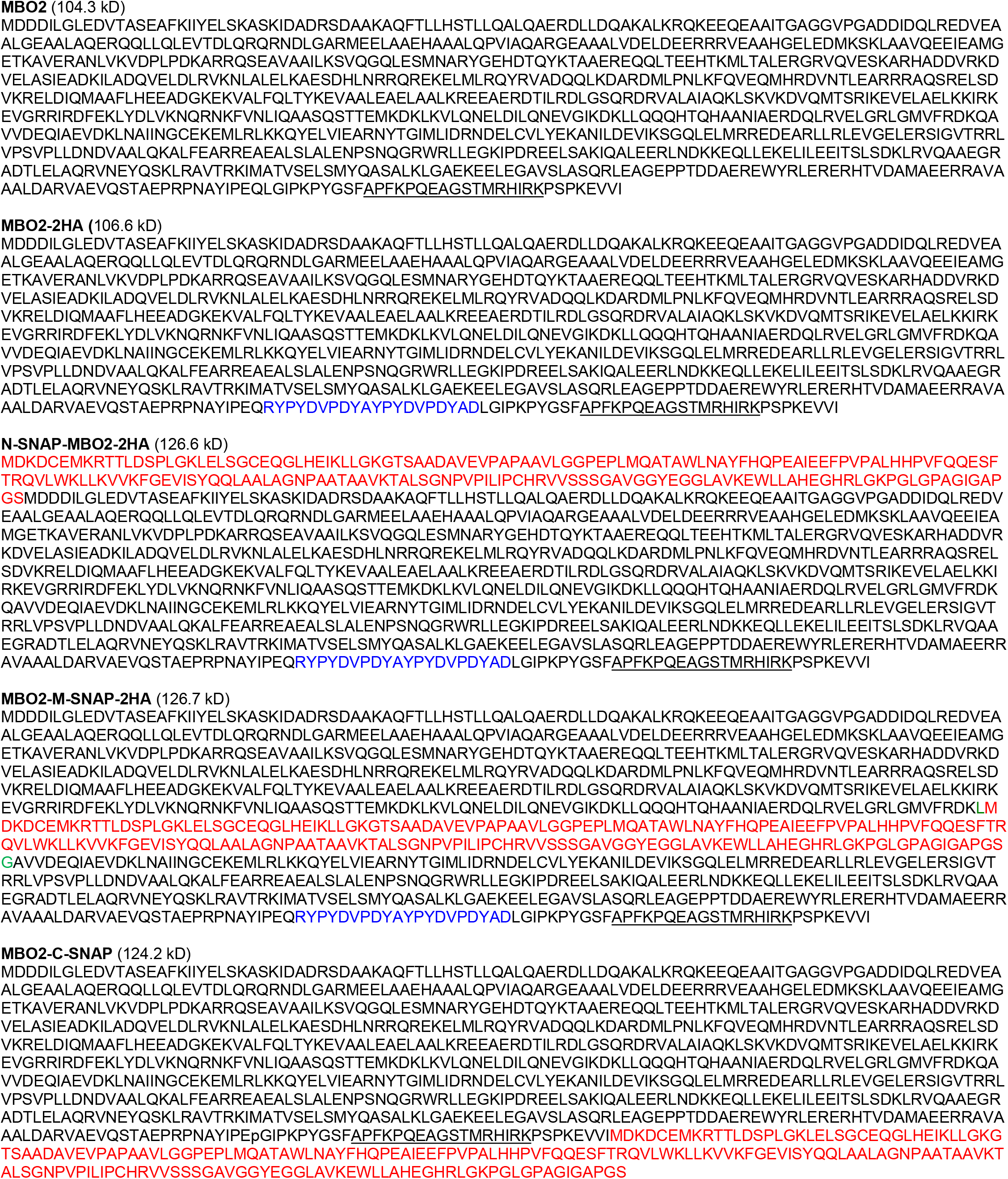
Predicted amino acid sequences of MBO2 and its tagged variants. The MBO2 sequence is shown in black, and the peptide sequence used for antibody production (Tam and Lefebvre, 2002) is underlined. The 2-HA-tag is shown in blue, and the SNAP-tags are shown in red.

**Figure S2.**
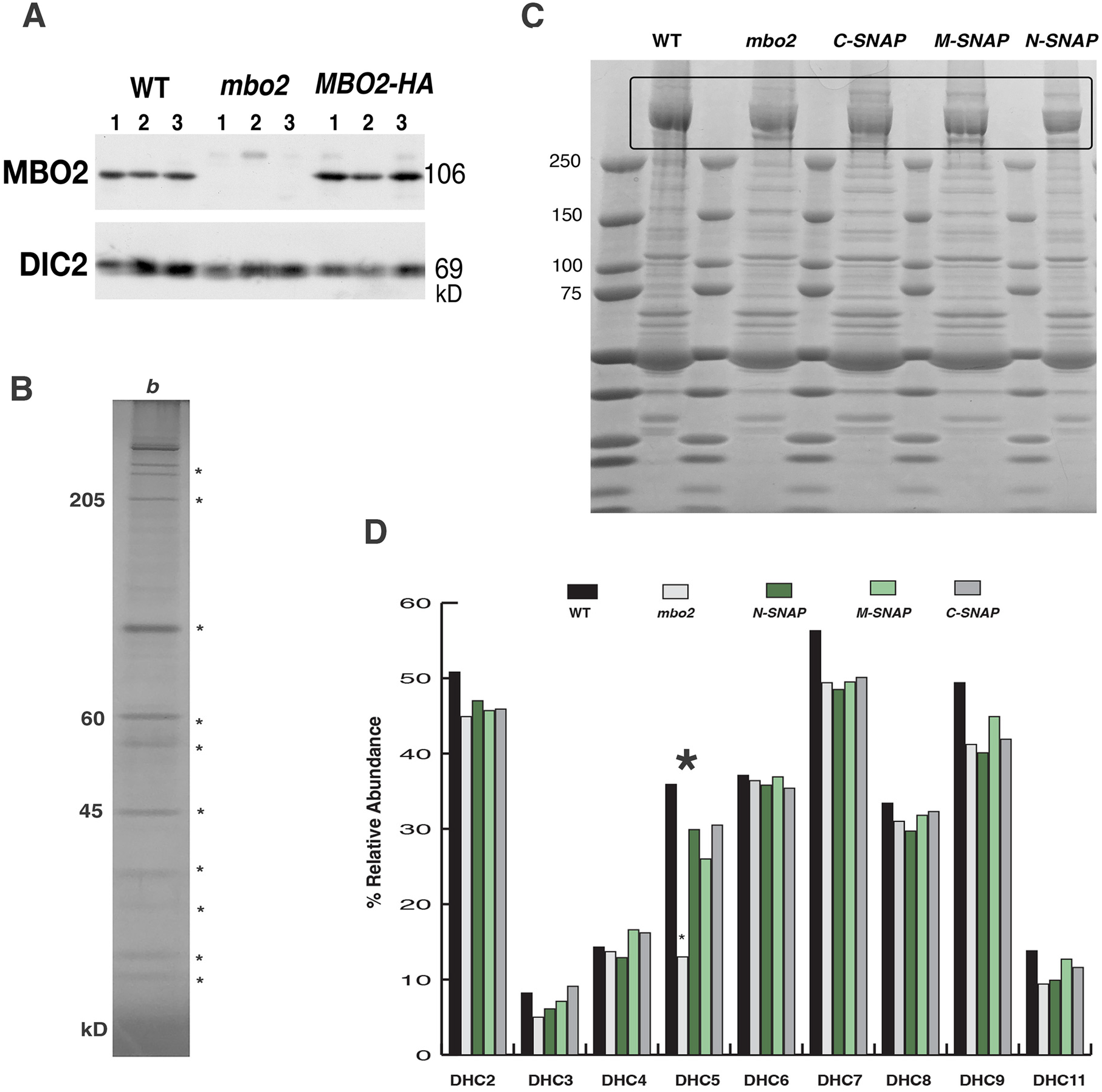
Protein samples used for MS/MS analysis and quantification of DHC content. **(A)** Western blot of three independent biological replicates of WT, *mbo2*, and *MBO2-HA* axonemes that were labeled using TMT isobaric tags and subjected to MS/MS. Blot was probed with antibodies to MBO2 and DIC2 (IC69). **(B)** A 0.6M extract of *pf28* was analyzed by FPLC and SDS-PAGE. The fraction containing the peak of IDA *b*/DHC5 was silver stained, and indicated bands were excised and analyzed by MS/MS. **(C)** Axonemes from WT, *mbo2*, and three SNAP-tagged rescued strains were analyzed by SDS-PAGE and stained with Coomassie Blue. The region containing the DHCs (outlined in black) was excised and analyzed by MS/MS. **(D)** The relative abundance of DHCs from the samples shown in (C) was determined by spectral counting. Only DHC5 was significantly reduced in *mbo2* (red arrowhead). DHC content was also quantified using the tools available in Proteome Discover (Supplemental Table S4).

**Figure S3.**
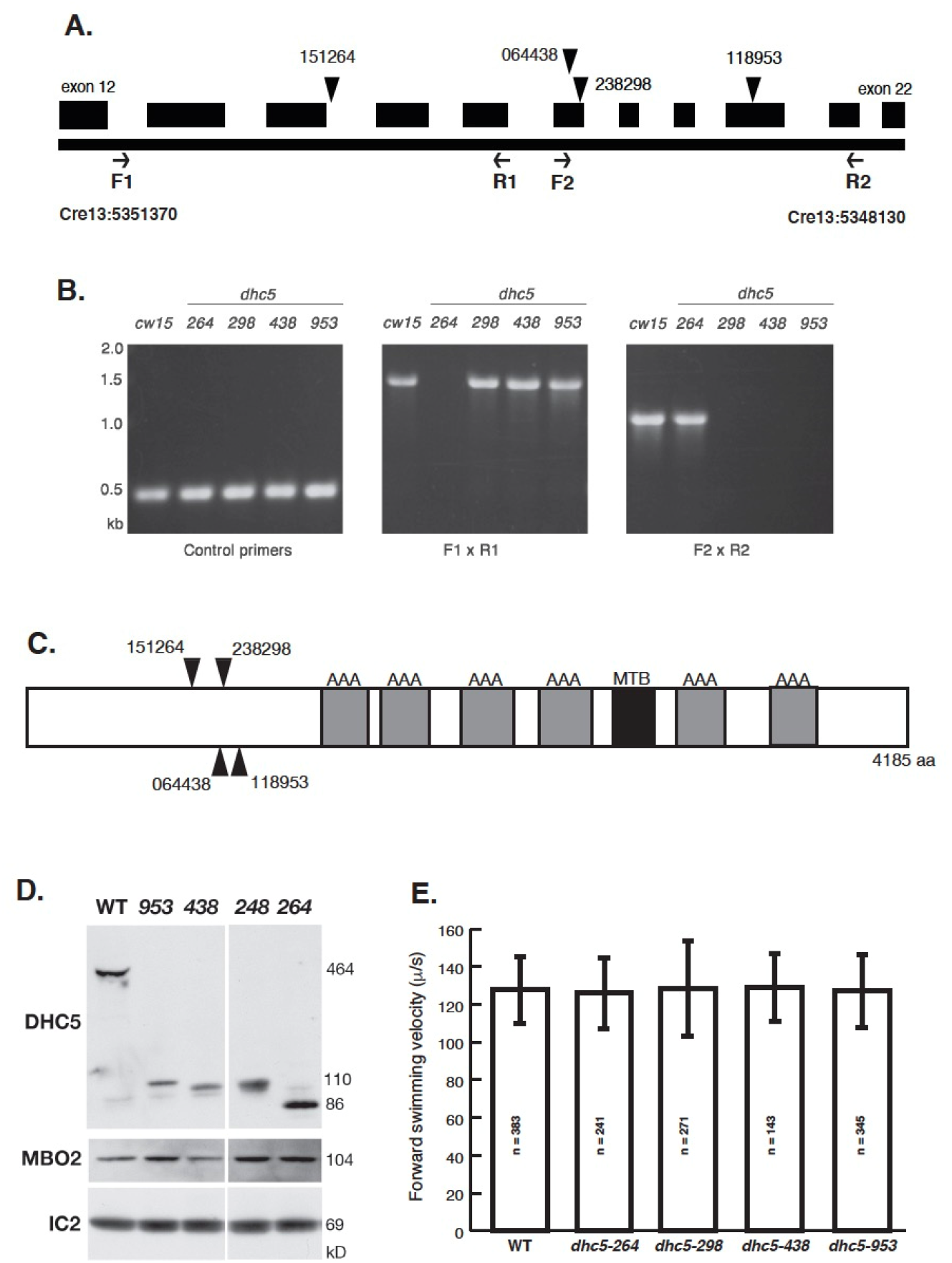
Loss of the DHC5 motor domain does not significantly alter forward swimming velocities. **(A)** Four *dhc5* strains generated by insertional mutagenesis were tested by PCR to confirm insertion sites in the 5’end of the *DHC5* gene encoding the N-terminal tail domain of DHC5. A small region of Chromosome 13 (Cre13) is shown here indicating the predicted plasmid insertion sites and the location of the PCR primers used to confirm the sites. **(B)** Genomic DNA was isolated from a control strain (*cw15*) and four *dhc5* candidates (referred to as *248, 264, 438, 953* based on the last 3 digits of the strain number) and subjected to PCR with control primers and experimental primers shown in (**A)** and Supplemental Table S2. **(C)** Schematic diagram of the DHC5 polypeptide showing the approximate location of the six AAA ATPase domains (gray), the microtubule binding domain (MTB, black), and the plasmid insertion sites in the N-terminal region (arrowheads). **(D)** Western blots of WT (*cw15*) and *dhc5* (*248, 264, 438, 953*) axonemes probed with different antibodies. N-terminal fragments of DHC5 lacking the motor domain were assembled in all four mutants. No significant defects were noted in assembly of MBO2 or DIC2 (IC69). **(E)** Measurements of forward swimming velocities failed to detect significant differences between WT and the four *dhc5* mutants.

**Figure S4.**
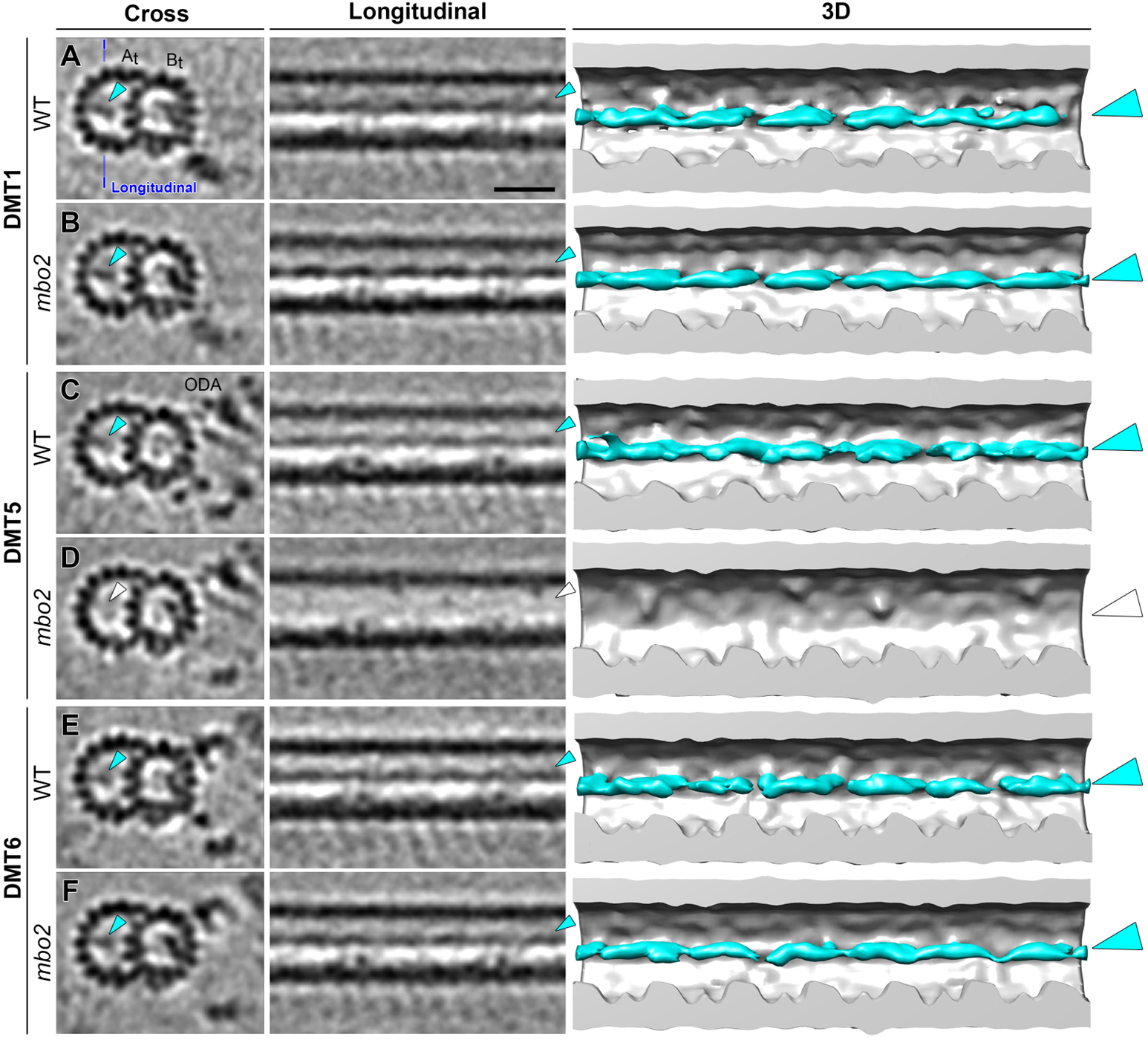
The B-tubule beak structure is missing in DMT5 in *mbo2*. **(A–F)** Tomographic slices (left two columns viewed in cross-sectional and longitudinal orientations) and 3D isosurface rendering (viewed in longitudinal orientation) show the B-tubule beak structure in DMT1 (**A** and **B**), DMT5 (**C** and **D**) and DMT6 (**E** and **F**) of WT and *mbo2* axonemes. The blue line in (**A)** indicates the location of the longitudinal slices. Cyan arrowheads indicate presence of the beak structure, which is missing in DMT5 in *mbo2* (**D**, white arrowheads). Scale bar in (A) is 20 nm (valid for all tomographic slices).

**Figure S5.**
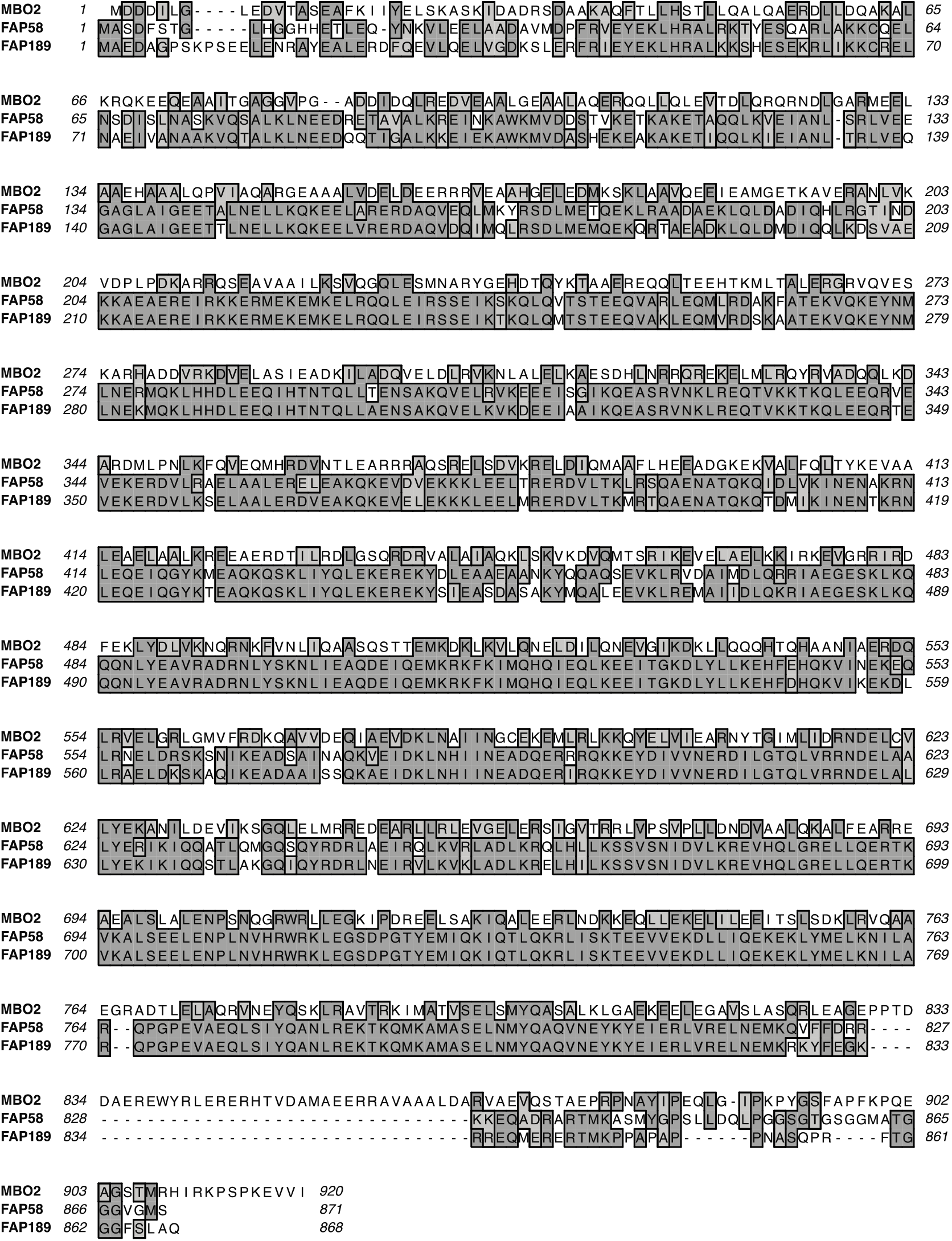
Clustal W alignment used for AlphaFold2 modeling of a predicted MBO2/FAP58 heterodimer. The full-length polypeptide sequences of MBO2, FAP58, and FAP189 were aligned by Clustal W, with identical amino acids shown in dark gray and similar amino acids in light gray.

**Figure S6.**
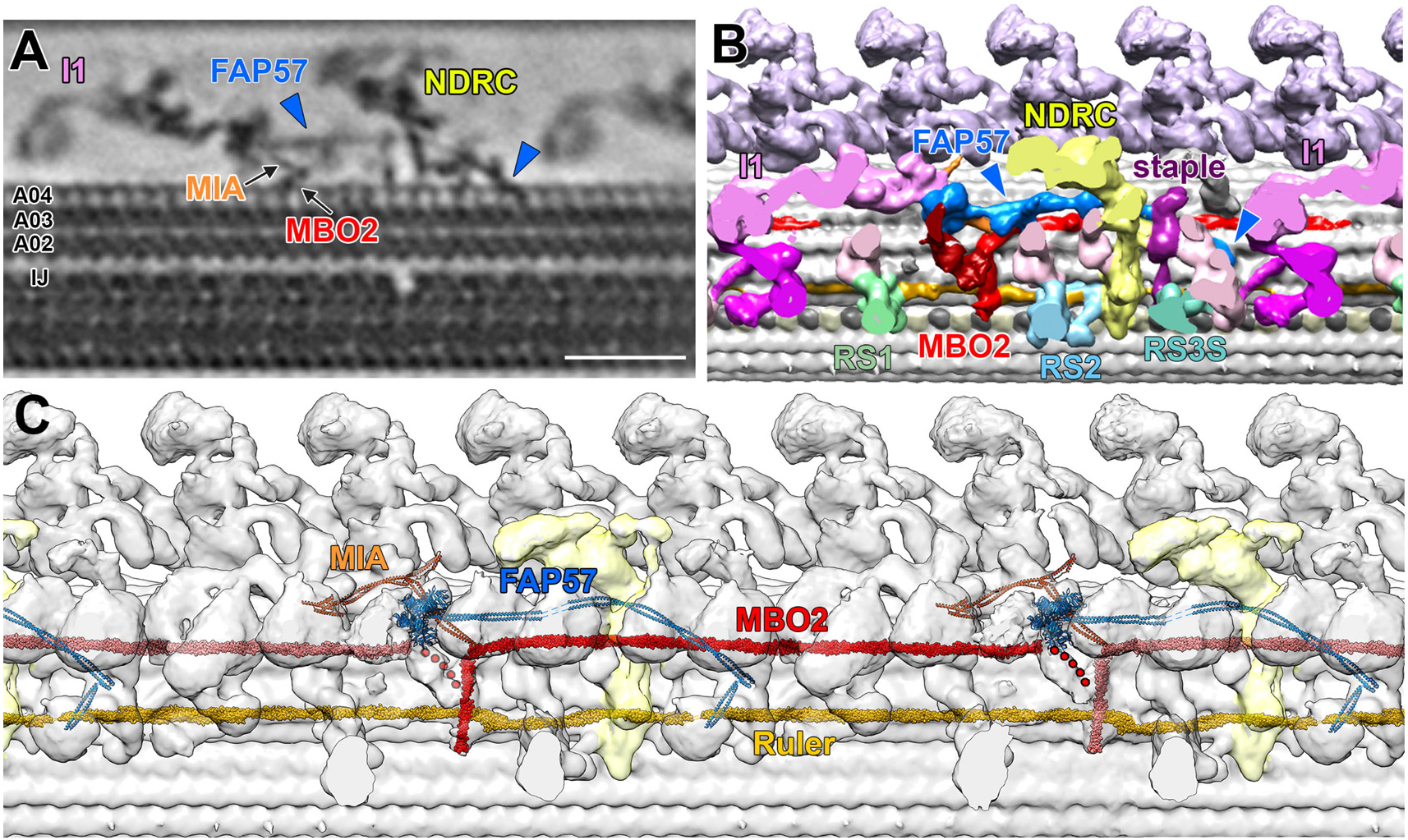
A model illustrating the proposed location of FAP57 relative to other structures in the 96 nm repeat. **(A)** Tomographic slice through an averaged 96-nm repeat from WT axonemes, showing the region projecting from the surface of the A-tubule above protofilament A04 and containing the ICLC complex of the I1 dynein and the N-DRC. A MIA filament extending from the base of the I1 ICLC is indicated in orange. The MIA complex contacts another filament on the surface of the DMT corresponding possibly to FAP189 and/or MBO2/FAP58. A second filament predicted to be the FAP57 coiled coil domain located above the surface of the DMT is indicated in blue. It extends from the base of a globular domain to a site beyond the N-DRC, curving down to a cleft between protofilaments A03 and A04 on the right (blue arrowheads). **(B)** Isosurface rendering of the structures shown in (A). The I1 dynein is sliced through one of its motor domains and colored in bright pink. The tether head structure below I1 is colored in magenta, and the bases of the other IDAs are colored in light pink. The CCDC39/40 complex forming the axonemal ruler between protofilaments A02 and A03 is shown in gold. The radial spokes (RS1, RS2, and RS3S) are cut close to their bases and colored in shades of green and light blue. The N-DRC structure is yellow, and structures missing in *mbo2* are colored in red. The blue-labeled structure corresponds to the proposed location of FAP57. Its N-terminal, WD repeat domain is located near the I1 ICLC and MIA complex (Yamamoto et al., 2013; Lin et al., 2019). Its coiled coil domain “floats” above the surface of the DMT, running behind the N-DRC and the “distal staple” (composed of CCDC96 and CCDC113, Bazan et al, 2021) and then curving down towards the bases of IDA *g* and IDA *d.* **(C)** A isosurface rendering of the WT 96 nm repeat with Alphafold2 predicted models of MIA complex (FAP100 and FAP73) and FAP57 (amino acids 1-771, 810-1050, 1060-1123) as predicted by Walton et al. (2023) shown relative to our model of the MBO2/FAP58 heterodimer. The C-terminal region of FAP57 (amino acids 1124-1316) is not shown. The FAP57 coiled coil domain is proposed to interact with the CCDC96/CCDC113 staple, FAP337, the base of IDA *g*, and the FAP43/44 subunits of the I1 tether head (Lin et al., 2019; Bazan et al., 2021, Ghanaeian et al., 2023; Walton et al., 2023). Note that *Chlamydomonas* contains three FAP57 paralogues (FAP57, FBB7, and FAP331) and two FAP337 paralogues (FAP337 and Cre07.g313850) (Lin et al., 2019); these three proteins may form homo- or heterodimers that differ between DMTs or proximal/distal regions of the axoneme. Scale bar is 25 nm in (A).

**Supplemental Table S1.**
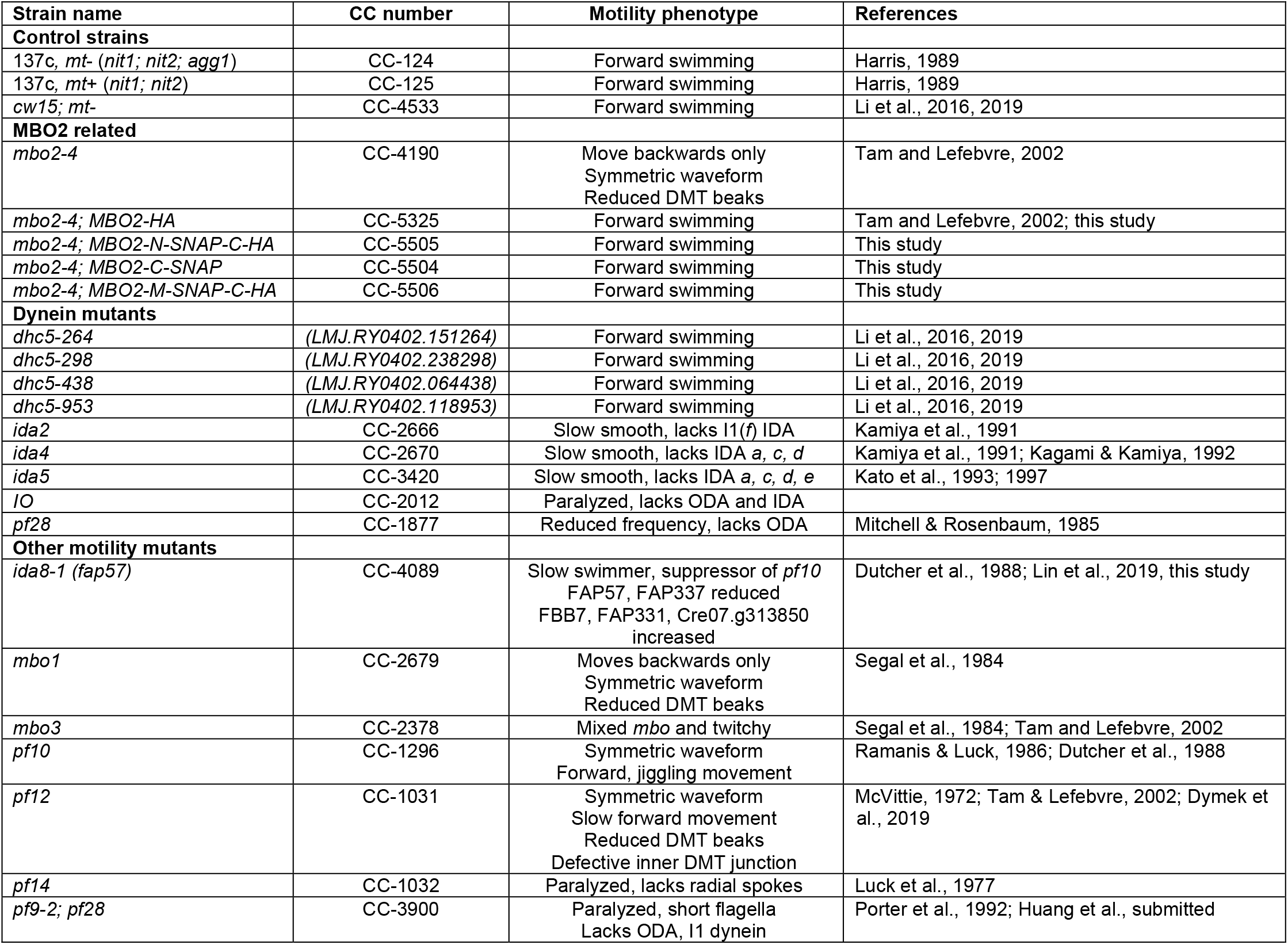
Strains used in this study.

| Strain name | CC number | Motility phenotype | References |
| --- | --- | --- | --- |
| **Control strains** |  |  |  |
| *137c, mt- (nit1; nit2; agg1)* | CC-124 | Forward swimming | Harris, 1989 |
| *137c, mt+ (nit1; nit2)* | CC-125 | Forward swimming | Harris, 1989 |
| *cw15; mt-* | CC-4533 | Forward swimming | Li et al., 2016, 2019 |
| **MBO2 related** |  |  |  |
| *mbo2-4* | CC-4190 | Move backwards only<br>Symmetric waveform<br>Reduced DMT beaks | Tam and Lefebvre, 2002 |
| *mbo2-4; MBO2-HA* | CC-5325 | Forward swimming | Tam and Lefebvre, 2002; this study |
| *mbo2-4; MBO2-N-SNAP-C-HA* | CC-5505 | Forward swimming | This study |
| *mbo2-4; MBO2-C-SNAP* | CC-5504 | Forward swimming | This study |
| *mbo2-4; MBO2-M-SNAP-C-HA* | CC-5506 | Forward swimming | This study |
| **Dynein mutants** |  |  |  |
| *dhc5-264* | (LMJ.RY0402.151264) | Forward swimming | Li et al., 2016, 2019 |
| *dhc5-298* | (LMJ.RY0402.238298) | Forward swimming | Li et al., 2016, 2019 |
| *dhc5-438* | (LMJ.RY0402.064438) | Forward swimming | Li et al., 2016, 2019 |
| *dhc5-953* | (LMJ.RY0402.118953) | Forward swimming | Li et al., 2016, 2019 |
| *ida2* | CC-2666 | Slow smooth, lacks I1(f) IDA | Kamiya et al., 1991 |
| *ida4* | CC-2670 | Slow smooth, lacks IDA *a*, *c*, *d* | Kamiya et al., 1991; Kagami & Kamiya, 1992 |
| *ida5* | CC-3420 | Slow smooth, lacks IDA *a*, *c*, *d*, *e* | Kato et al., 1993; 1997 |
| *IO* | CC-2012 | Paralyzed, lacks ODA and IDA |  |
| *pf28* | CC-1877 | Reduced frequency, lacks ODA | Mitchell & Rosenbaum, 1985 |
| **Other motility mutants** |  |  |  |
| *ida8-1 (fap57)* | CC-4089 | Slow swimmer, suppressor of *pf10*<br>FAP57, FAP337 reduced<br>FBB7, FAP331, Cre07.g313850<br>increased | Dutcher et al., 1988; Lin et al., 2019, this study |
| *mbo1* | CC-2679 | Moves backwards only<br>Symmetric waveform<br>Reduced DMT beaks | Segal et al., 1984 |
| *mbo3* | CC-2378 | Mixed *mbo* and twitchy | Segal et al., 1984; Tam and Lefebvre, 2002 |
| *pf10* | CC-1296 | Symmetric waveform<br>Forward, jiggling movement | Ramanis & Luck, 1986; Dutcher et al., 1988 |
| *pf12* | CC-1031 | Symmetric waveform<br>Slow forward movement<br>Reduced DMT beaks<br>Defective inner DMT junction | McVittie, 1972; Tam & Lefebvre, 2002; Dymek et al., 2019 |
| *pf14* | CC-1032 | Paralyzed, lacks radial spokes | Luck et al., 1977 |
| *pf9-2; pf28* | CC-3900 | Paralyzed, short flagella<br>Lacks ODA, I1 dynein | Porter et al., 1992; Huang et al., submitted |

**Supplemental Table S2.**
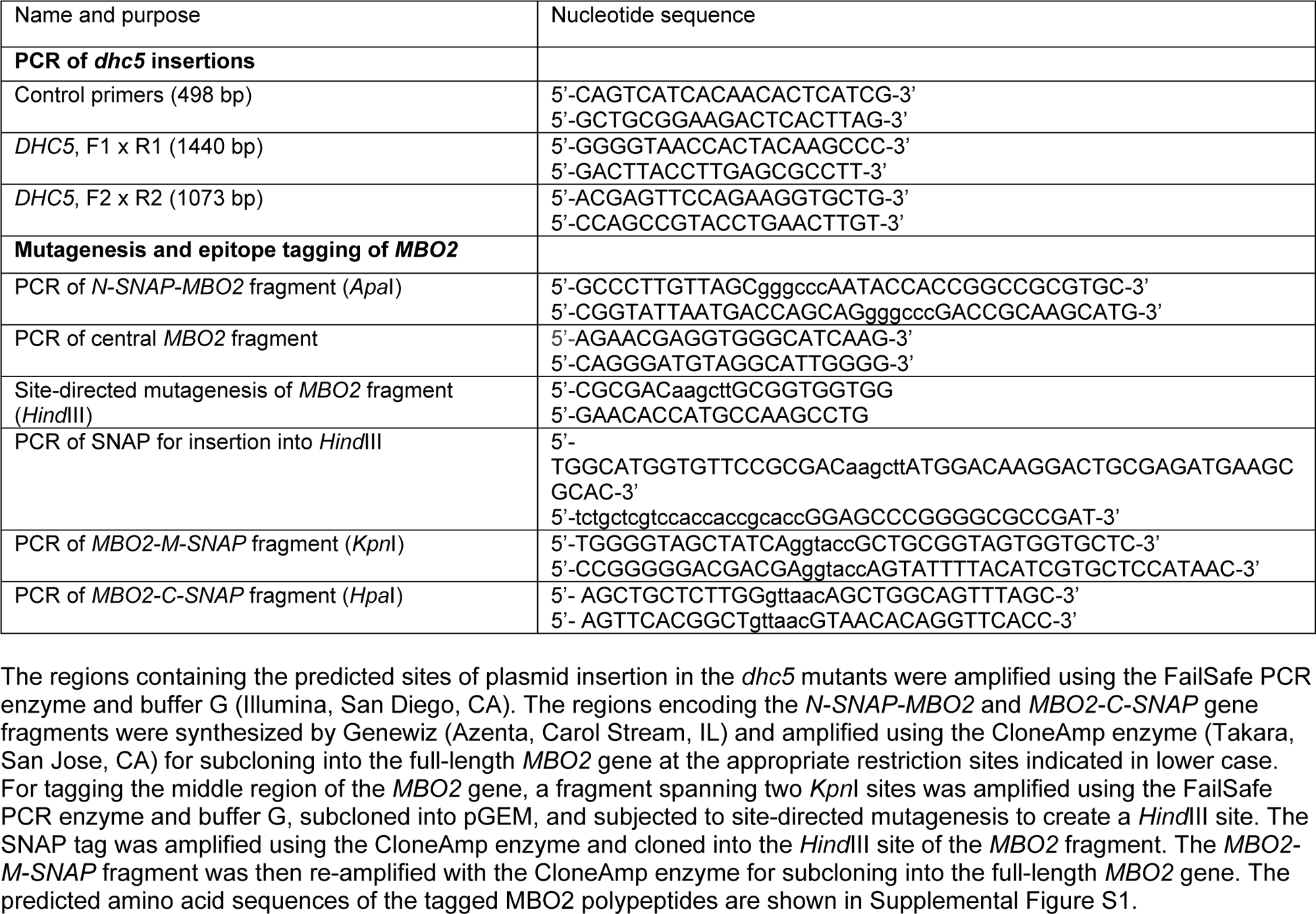
Oligonucleotide sequences used in the study.

**Supplemental Table S3.** Antibodies used in this study.

| Antigen | Host | Dilution (WB) | Reference or source |
| --- | --- | --- | --- |
| DRC1 | Rabbit | 1:1000-1:10000 | Wirschell et al., 2013 |
| Gas8 fusion (DRC4) | Rabbit | 1:1000-1:10000 | Bower et al., 2013 |
| CCDC39/FAP59 | Rabbit | 1:1000 | Sigma #HPA035364 |
| FAP57 | Rabbit | 1:1000-5000 | This study |
| Rib43 | Rabbit | 1:10000-1:20000 | Norrander et al., 2000 |
| Rib72 | Rabbit | 1:10000-1:20000 | Ikeda et al., 2003 |
| tektin | Rabbit | 1:20000 | Yanagisawa & Kamiya, 2004 |
| DHC5 | Rabbit | 1:1000 | Yagi et al., 2009 |
| DHC9 | Rabbit | 1:5000 | Yagi et al., 2009 |
| IC140 | Rabbit | 1:10000 | Yang & Sale, 1998 |
| IC2 (IC69) | Mouse | 1:10000-1:20000 | Sigma #D6168 |
| IC138 | Rabbit | 1:10000 | Hendrickson et al., 2004 |
| MIA1 | Rabbit | 1:1000 | Yamamoto et al., 2012 |
| MIA2 | Rabbit | 1:1000 | Yamamoto et al., 2012 |
| MBO2 | Rabbit | 1:1000 | Tam and Lefebvre, 2002 |
| RSP16 | Rabbit | 1:10000-1:20000 | Yang et al., 2005 |
| HA (3F10) | Rat | 1:500-1000 | Roche #1867423 |
| GFP | Mouse | 1:5000 | Covance #MMS-118P |
| SNAP | Rabbit | 1:1000 | New England Biolabs #P9310S |

**Supplemental Table S5.** Summary of image processing information for strains used in this study.

| Strains | Tomogram numbers<br>(proximal/distal) | Averaged<br>DMT repeats | TEM/<br>camera | Resolution<br>(nm)<sup>a</sup> | Usage in Figures |
| --- | --- | --- | --- | --- | --- |
| WT | 25 (5/20) | 3736 | F30/CCD | 3.8 | Figure 2A-D, I-K<br>Figure 3, Figure 4E, F<br>Figure 5F,<br>Figure S5A, C, E |
| WT | 19 (6/13) | 2381 | Krios/K2 | 2.3 | Figure 5A, C,<br>Figure 6A, B, |
| pWT<sup>b</sup> | 85 (26/59) | 11519 | Krios/K2 | 1.8 | Figure 7, Figure S7 |
| *mbo2* | 37 (10/27) | 4525 | F30/CCD | 3.1 | Figure 2E-H, I-K<br>Figure 3, Figure 4E, F<br>Figure S5B, D, F |
| *mbo2; MBO2::SNAP* (C) | 14 (4/10) | 1898 | F30/CCD | 3.7 | Figure 4E, F, Figure 5F |
| *mbo2; SNAP::MBO2* (N) | 14 (5/9) | 2257 | Krios/K2 | 2.4 | Figure 4E, F, Figure 5B |
| *mbo2; MBO2::SNAP* (M) | 25 (7/18) | 3154 | Krios/K2 | 2.5 | Fig4E, F, Figure 5D |

<sup>a</sup> Resolution based on the 0.5 criterion of the Fourier shell correlation
<sup>b</sup> Strains include WT, *fap76-1*, *fap81*, *fap92* and *fap76; fap81*, which were used to analyze central apparatus. The central apparatus mutants had WT-like 96-nm axonemal DMT repeats (Fu et al., 2019).

